# Thalamic encoding of complex sensory patterns and its possible role in cognition

**DOI:** 10.1101/2020.08.19.257667

**Authors:** Carlos Castejon, Jesus Martin-Cortecero, Angel Nuñez

## Abstract

The function of the higher-order sensory thalamus remains unresolved. Here, POm nucleus was examined by *in vivo* extracellular recordings across a range of complex sensory patterns. We found that POm was highly sensitive to multiwhisker stimuli involving complex spatiotemporal interactions. The dynamical spatiotemporal structure of sensory patterns and the different complexity of their parts were accurately reflected in precise POm activity changes. Importantly, POm was also able to respond to ipsilateral stimulation and was implicated in the representation of bilateral tactile events by integrating simultaneous signals arising from both whisker pads. We found that POm nuclei are mutually connected through the cortex forming a functional POm-POm loop. We unravelled the nature and content of the messages travelling through this loop showing that they were ‘structured patterns of sustained activity’. These structured messages were transmitted preserving their integrated structure. The implication of different cortical areas was investigated revealing that S1 plays a protagonist role in this functional loop. Our results also demonstrated different laminar implication in the processing of sustained activity in this cortical area and its transmission between hemispheres. We propose a theoretical model in which these ‘structured patterns of sustained activity’ generated by POm may play important roles in perceptual, motor and cognitive functions. From a functional perspective, this proposal, supported by the results described here, provides a novel theoretical framework to understand the implication of the thalamus in cognition. In addition, a profound difference was found between VPM and POm functioning. The hypothesis of Complementary Components is proposed here to explain it.

**Highlights:**

POm is implicated in the representation of complex sensory patterns.
POm is implicated in the encoding of bilateral tactile events.
POm nuclei are mutually connected through the cortex forming a functional POm-POm loop.
‘Structured patterns of sustained activity’ travelling through the loop

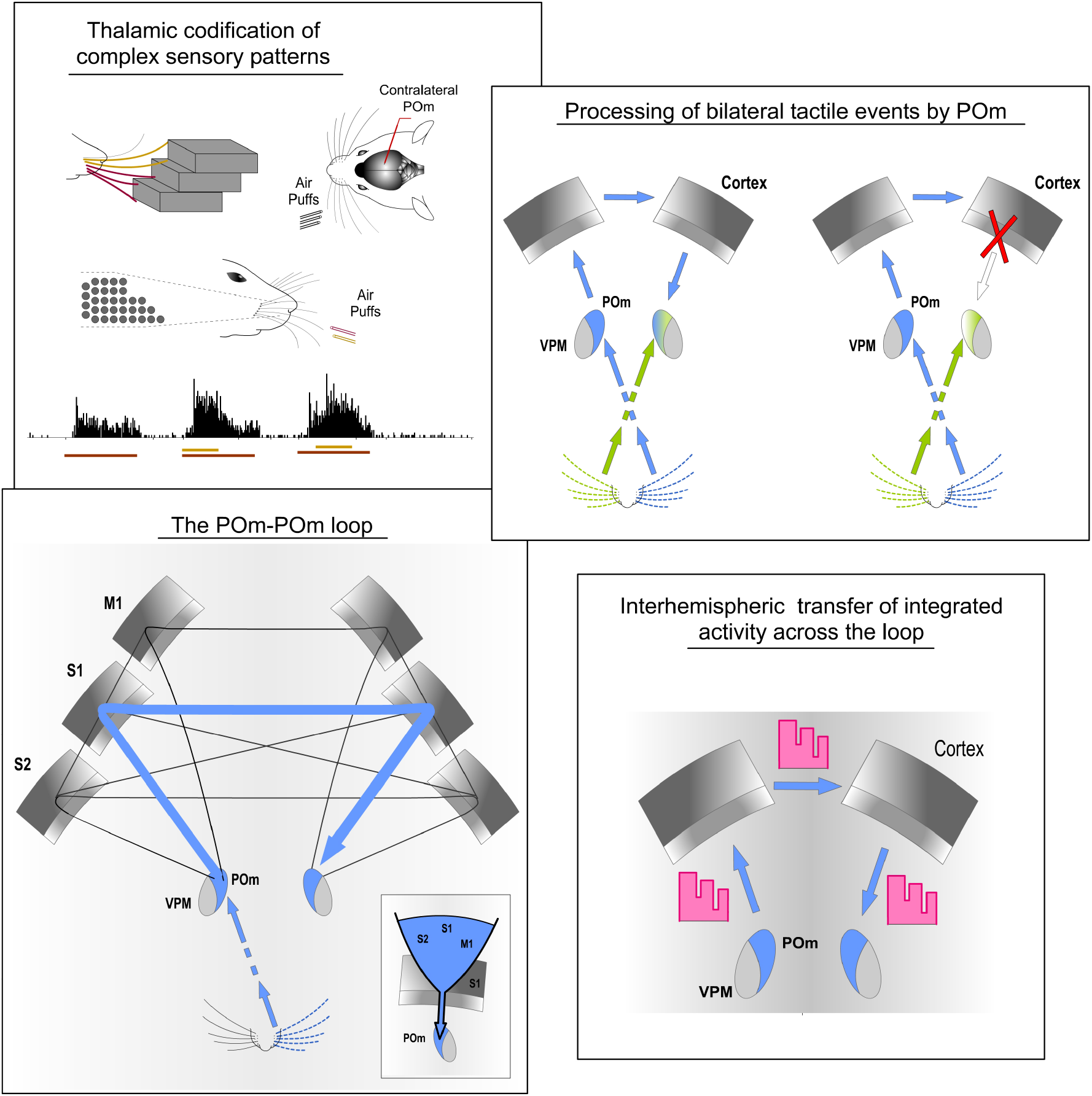

## Introduction

Traditionally, sensory systems have mostly been studied using simple and discrete stimuli. However, in natural conditions, sensory events usually have complex and dynamical spatiotemporal structures and normally multiple sensory signals occur simultaneously with different onsets, offsets and overlappings, challenging the computational capacities of sensory systems. However, it is still unclear how sensory systems generate a representation of these dynamics. Moreover, how sensory systems extract relevant patterns from the raw sensory flow is poorly understood. Here, we propose the hypothesis that higher-order sensory thalamus has a protagonist role in that function.

The rodent whisker system has an extraordinary ability to extract patterns and regularities from the environment and provides a perfect model in which to test our proposal. Rodents have an array of whiskers on each side of the face and during tactile exploration, multiple whiskers are stimulated simultaneously. Accordingly, the activation of individual whiskers strongly overlaps. These multiple contacts with the whiskers generate complex patterns of sensory information. How the somatosensory system transforms these merged raw sensory signals into reliable neural representations and extracts information from that apparent noise is still unclear.

In these animals, tactile information from whiskers is processed by two main parallel ascending pathways towards the cortex (Diamond et al., 1992; Veinante et al., 2000a; Ahissar et al. 2000). The lemniscal pathway includes the principal trigeminal nucleus (Pr5) and the ventral posteromedial thalamic nucleus (VPM). The paralemniscal pathway includes the spinal trigeminal subnucleus interpolaris (Sp5i) and the posteromedial thalamic nucleus (POm). Although the function of VPM has been broadly studied, less is known about the function of POm. POm is powerfully driven by simultaneous activation of multiple whiskers. However, the functional implication of this characteristic remains unknown. Moreover, the content of POm representations and the nature of the messages that POm transfers to and receives from the cortex remain unclear.

In addition, sensory events are usually characterized by complex bilateral sensory patterns. Therefore, the integration of tactile information from the two sides of the body seems to be fundamental in the encoding of sensory patterns in bilateral perceptual function. Although, somatosensory cortical implication in the processing of bilateral stimuli has been much more studied, the implication of the thalamus in these tactile interactions remains unknown.

The following experiments were thought to study the implication of POm in the encoding of these complex phenomena.

## Results

### 1. Sustained activity in POm

Whisker-evoked responses in POm were examined by *in vivo* extracellular recordings using stimuli with different durations within the range used by these animals during their natural explorations (Fig. 1 and 2). While studying multi-unit activity (n = 119), we found that this nucleus was able to generate sustained responses even to long-duration stimuli. This capacity was consistent across all animals (n = 12). The duration of the stimulus did not alter the mean onset latency of responses (p = 0.52, One-way ANOVA; Fig. 2B).

**Fig. 1.**
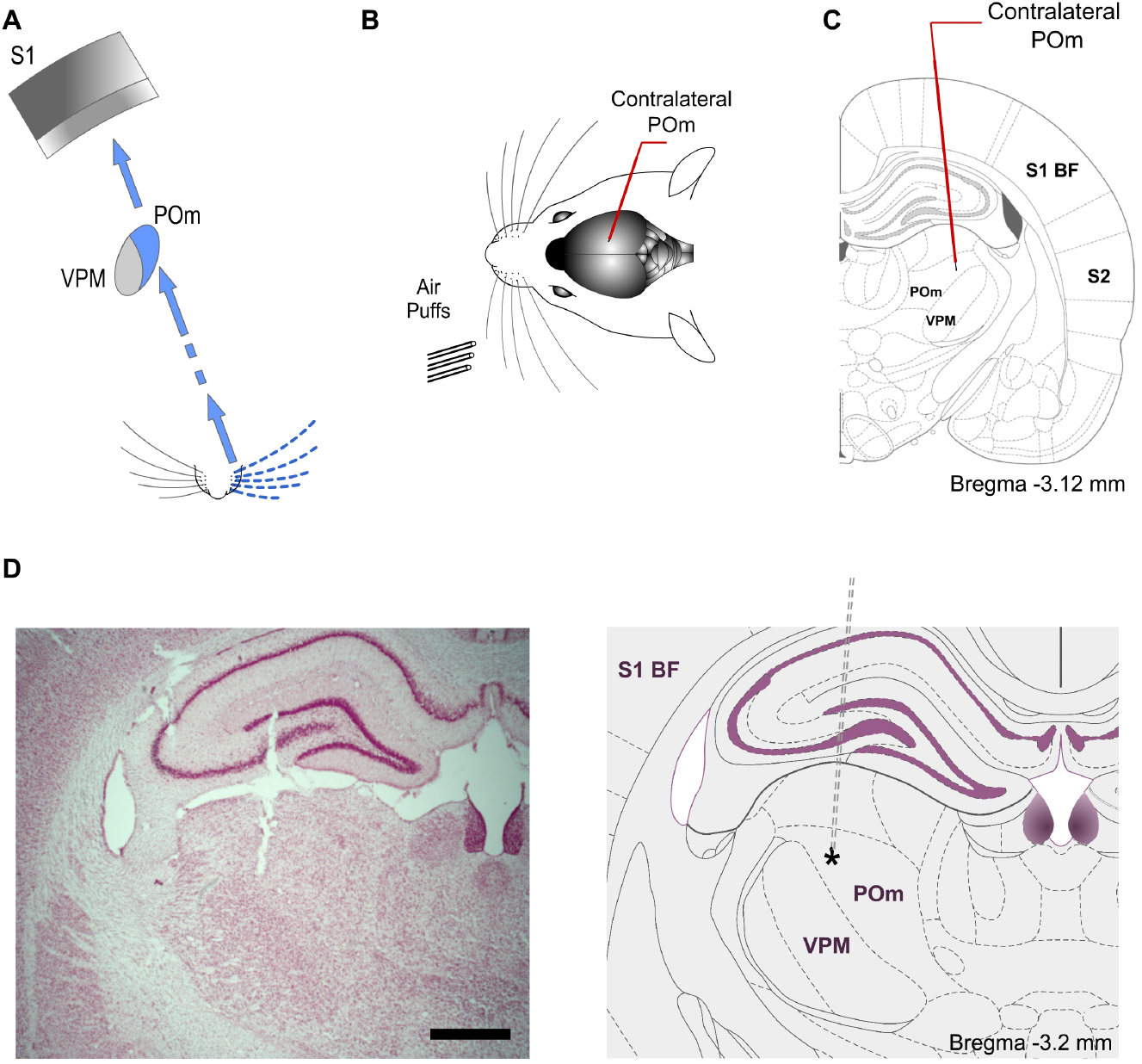
Experimental paradigm. (A) Illustration of the paralemniscal pathway. (B) Schematic drawing displaying the sensory stimulation via patterns of multiwhisker deflections. Recordings were made in the contralateral POm (C). Coronal section illustrating a recording electrode inserted into POm. Bregma anteroposterior level is indicated. (D) The left panel shows a representative Nissl stained coronal section displaying the location of the recording site in the dorsolateral part of POm and the track left by the electrode. An atlas schematic reconstruction of this recording site within POm is shown in the right panel (Paxinos and Watson 2007). Tip position is indicated by an asterisk. Scale bar, 1 mm. S1 BF, primary somatosensory cortex barrel field. S2, secondary somatosensory cortex.

**Fig. 2.**
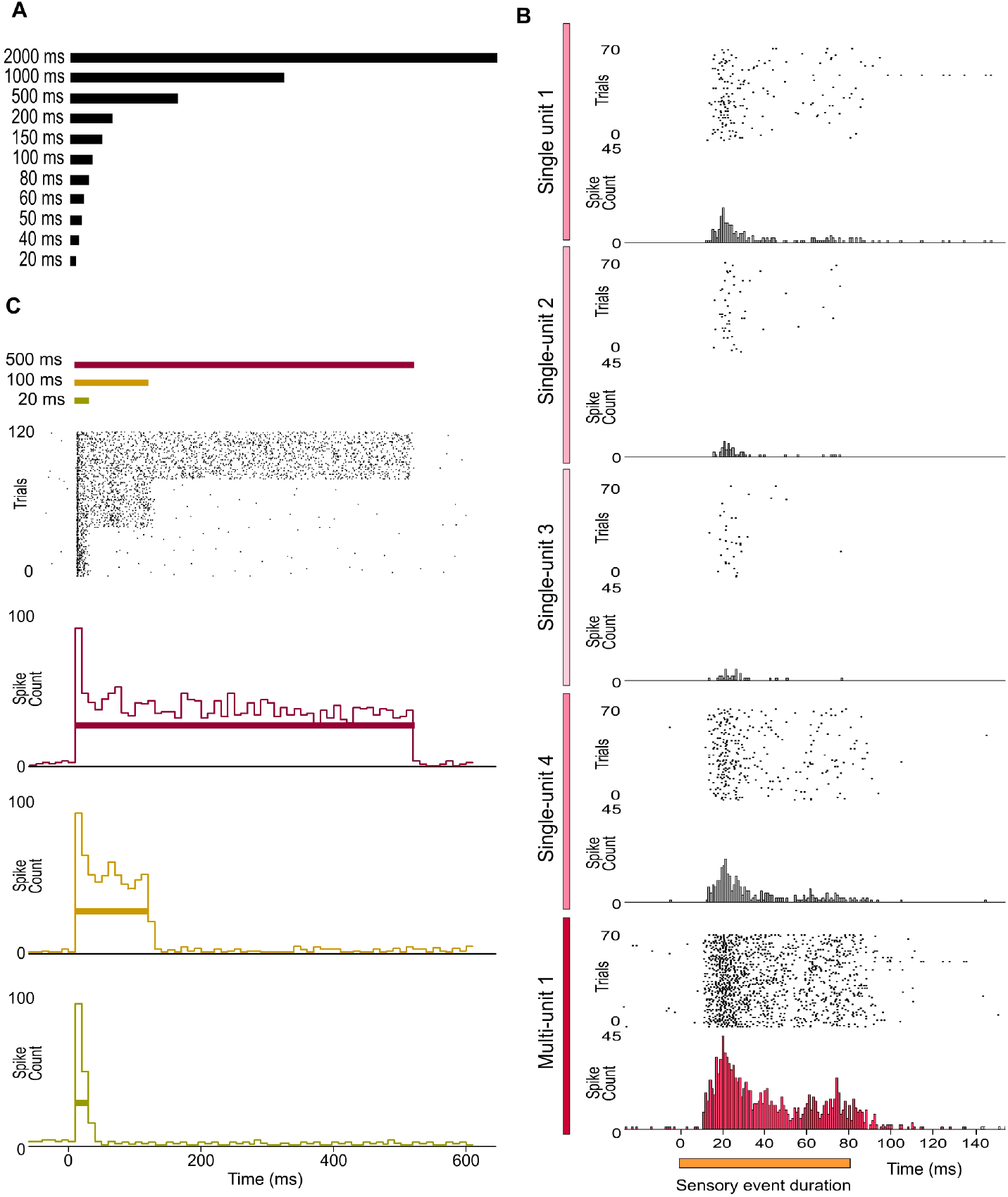
Sustained activity in POm. (A) Air-puffs used for sensory stimulation varied in duration from 20 ms to 2 seconds. (B) The encoding of the duration of sensory events by POm was studied at single-unit and multi-unit level. Raster plots and PSTHs showing one multi-unit and four single-units POm responses extracted from the same recording and evoked by a sensory pattern of 80 ms duration. Note that the robustness of this form of encoding by POm sustained activity is obtained from population response formed by the superposition of spikes from individual neurons. The gradual color intensity of vertical lines represents a simulated contribution of each single-unit in this example to the encoding of stimulus duration. (C) POm responds throughout the entire duration of the stimuli. Raster plot and peristimulus time histograms (PSTHs; bin width 10 ms) showing sustained multi-unit POm responses evoked by different stimulus duration (40 trials shown for each stimulus). Color lines indicate the duration of the stimulus. Time 0 indicates the onset of the stimulus. Note that the duration of the stimulus did not alter the onset latency of responses.

To analyse how this multi-unit activity replicated the response profile of individual POm neurons, we also characterized this phenomenon studying POm single-units (n = 91). We found that POm neurons responded homogeneously tending to generate sustained responses. This indicated that the population response formed by the combination of spikes from these neurons allowed for the encoding of stimuli duration by POm sustained activity (Fig. 2C). Thus, we mainly used POm multi-unit responses for further analyses.

Confirming previous findings (Castejon et al. 2016), these results show that POm has the capacity to sustain its activity to encode and represent tactile input duration with high accuracy. Given that POm can generate sustained responses greatly outlasting the duration of multiple whisk cycles, this capacity can be used to codify sensory patterns and sequences of stimuli. The following experiments were designed to study the implication of POm in the encoding of these phenomena.

### 2. POm encoding of spatiotemporal sensory patterns

During whisking rats integrate signals from many whiskers to obtain accurate tactile information from their environment. Our next experiments were designed to map how complex sensory information, produced when multiple whiskers are activated simultaneously during a tactile event, is encoded in the activity of POm.

#### 2.1. POm spatial integration of multiwhisker stimulation

First, we characterized the response properties of POm to simple spatial overlapping stimuli precisely delivering simultaneous air-puffs to different whiskers at different locations across the whisker pad. Consistent with published data (Diamond et al. 1992; Ahissar et al. 2000), we found large multiwhisker receptive fields (mean receptive field size: 10.9 ± 3.1 whiskers; n = 42 units). We observed that POm responses exhibited sustained increases in firing rate when different whiskers were activated simultaneously. To investigate this POm capacity in detail, we studied the integration of signals by the POm across a range of reproducible spatial overlapping combinations (Fig. 3A). Across them, we found that POm multi-unit responses showed sustained increases, as reflected in response magnitude, during overlappings of spatial signals (quantified in Fig. 3C). The spatial integration was also observed between remote whiskers. The mean onset latency and duration of POm responses to different signals were not changed by their overlapping (Fig 3C). We also investigated these effects studying POm single-units (quantified in Fig. 3B). Their responses showed similar dynamics to multi-unit responses. These findings were consistent across all animals (n = 15). Importantly, the robustness of this capacity of encoding seems to be obtained from population response (Fig. 3C). Accordingly, we used POm multi-unit responses for further analyses of POm integration.

**Fig. 3.**
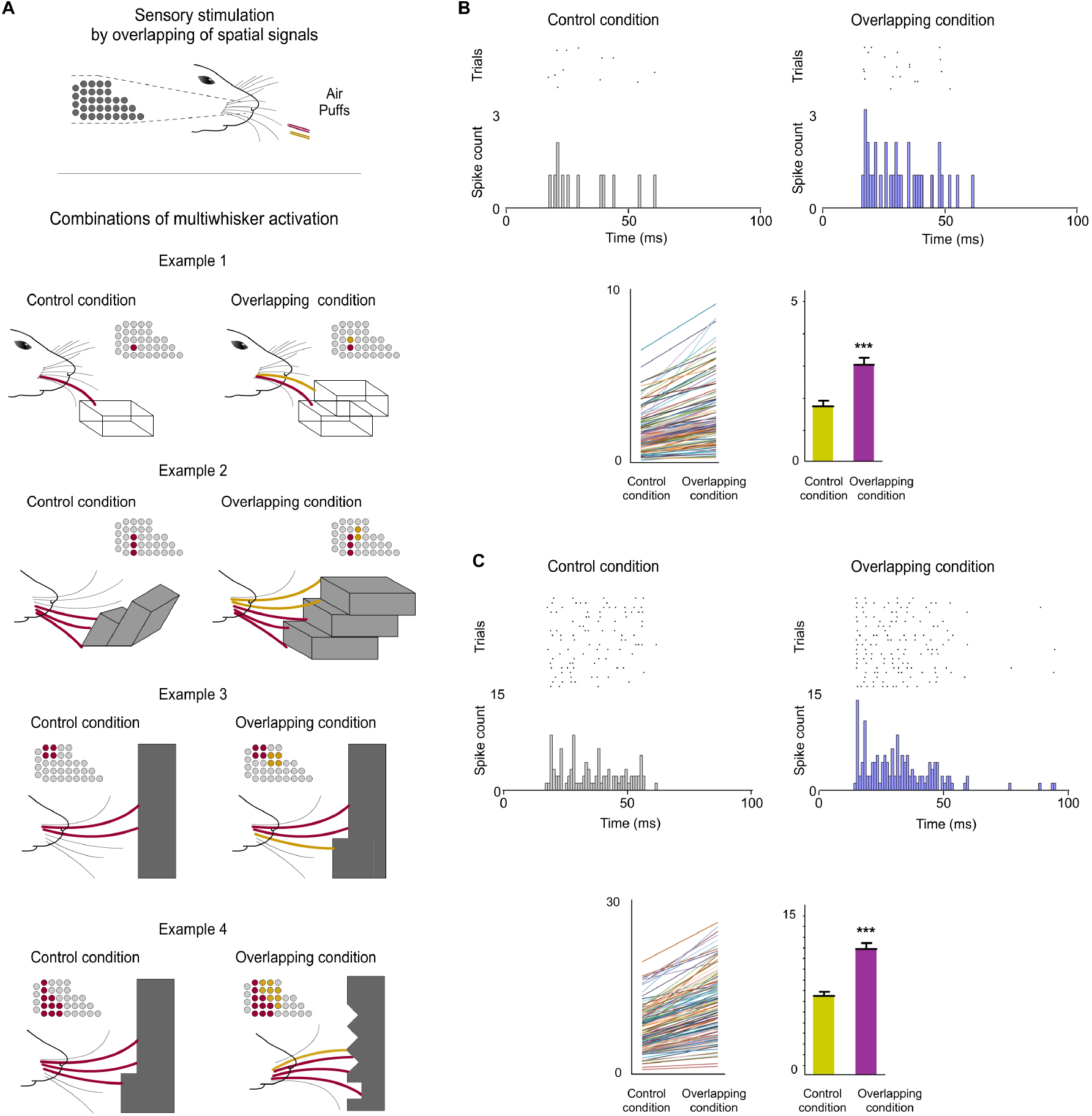
POm integration of spatial signals. (A) Sensory stimulation was produced by the application of individual (Control Condition) or simulataneous air puffs (Overlapping Condition) activating different whiskers and producing diverse combinations of multiwhisker activation. To simulate natural stimuli, overlappings produced by the activation of whiskers in different directions were also included. The number of whiskers activated by each air puff was varied to generate different combinations. They were repeated multiple times (range 20–70 repeats) and POm responses to them were studied and quantified by comparison between Control and Overlapping conditions. Four examples of these combinations, the whiskers activated (depicted in different colors in the schematic representations of the whisker pads) and their corresponding illustrations of their simulated possible real occurrence during the exploration of different objects, surfaces and textures in natural conditions are shown. (B) POm response magnitude increased when whiskers were activated simultaneously. This facilitative integration during overlappings of spatial signals showed that POm was activated more strongly by complex patterns than by simple ones as can be appreciated in the peri-stimulus time raster plots and histograms of a representative POm single-unit response for 20 trials evoked by the example 2 in A. Note the significant increase in the spike count during Overlapping condition. Data comparing the spike rate in Control and Overlapping conditions of single-units (n = 102, depicted in different colors) across stimulation combinations and the total mean response magnitude in both conditions are also shown. The mean firing rate of all single-units was significantly increased in the Overlapiing condition (72 %; p < 0.001; Wilcoxon matched-pairs test). (C) Same as in B, but for multi-unit activity. Raster plots and PSTHs of a demonstrative POm multi-unit response for 40 trials evoked by the example 3 in A. The spike rate in Control and Overlapping conditions of multi-units (n = 136, depicted in different colors) across stimulation combinations and the total mean response magnitude in both conditions are shown. The mean firing rate of all multi-units was significantly increased in the Overlapiing condition (56 %; p < 0.001; Wilcoxon matched-pairs test) Note that response duration was not altered by the overlapping.

Together, these results showed that POm was activated more strongly by complex stimuli than by simple ones. This was produced by a facilitative integration of overlapping spatial signals by POm.

#### 2.2. POm integration of spatiotemporal overlapping dynamics

Next, since tactile events typically have spatiotemporal structures that change dynamically in time, we studied how spatial integration (spatial dimension) occurs throughout the duration of the sensory pattern (temporal dimension) in 18 rats (Fig. 4). Interestingly, when overlappings of spatial signals were produced by delivering simultaneous air-puffs with different durations or with the same duration but applied at different times (Fig. 4A middle panel), increases in sustained responses were only observed during the overlapping time between them (data showing the quantification of this effect are described in Fig. 4D). The spatial integration of multiwhisker activation was only produced during the temporal overlapping. Importantly, the increases of POm activity were sustained along the temporal overlappings (Fig.4C). Therefore, the time shared by overlapping signals is also encoded by POm activity.

**Fig. 4.**
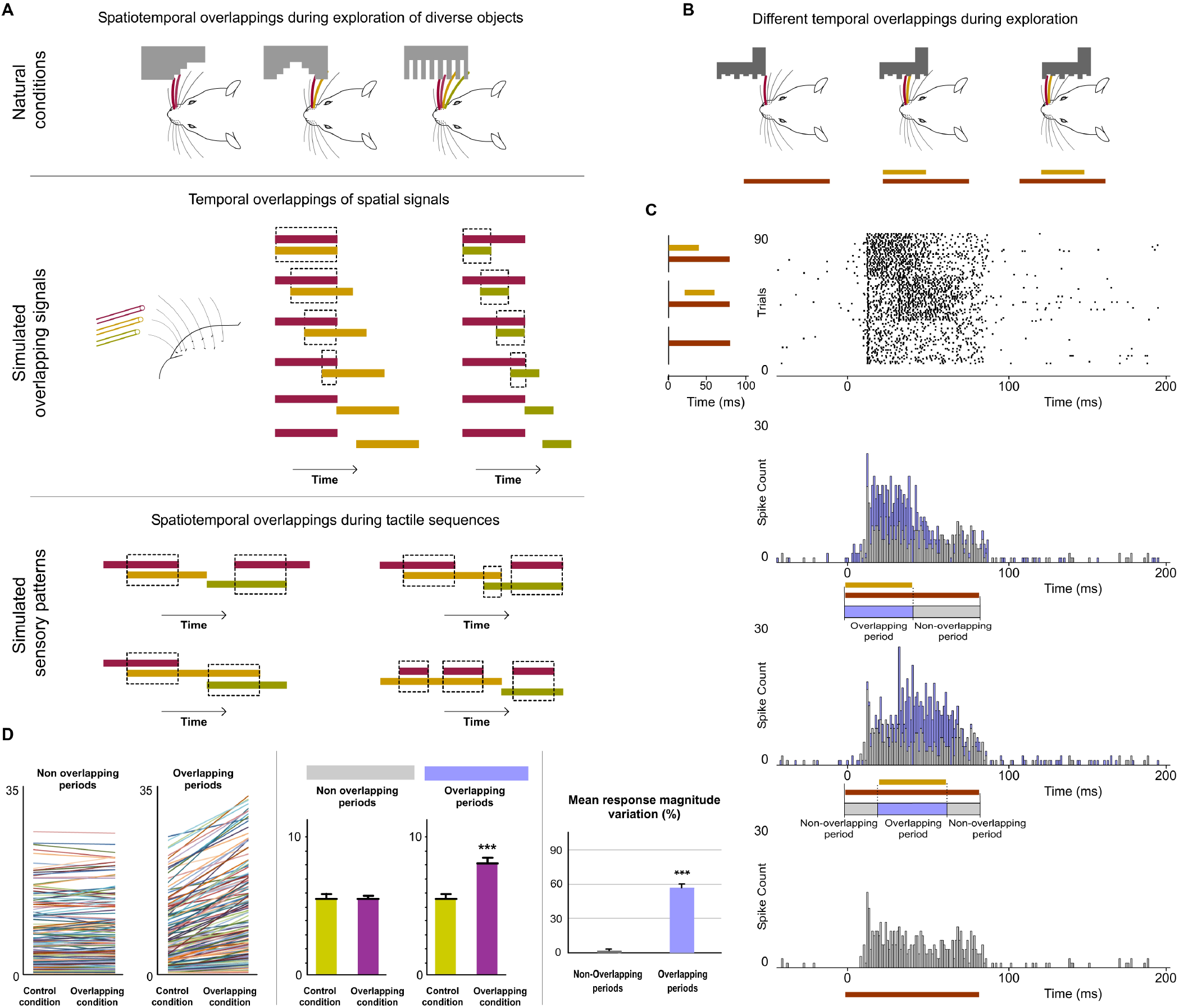
POm integration of spatiotemporal overlapping dynamics. (A) In natural conditions, diverse shapes and textures generate different sequences of multiwhisker activations and different dynamical overlappings between them during their exploration. Natural sensory patterns can be simulated by simultaneous activation of different whiskers and by varying the number of whiskers activated, their combination, order and duration of air pulses to produce different spatiotemporal overlappings. A variety of spatiotemporal patterns were designed to simulate possible real complex stimuli or sequences of stimuli using controlled multiwhisker deflections performed by overlapping air puffs of different durations applied to different whiskers including neighbouring whiskers or whiskers farther apart across the whisker pad. The duration of air puffs and the temporal overlapping between them were varied to generate different combinations (protocols). The number of whiskers activated by each air puff was varied to generate different sensory patterns using the same protocol. Short duration (20 ms) stimuli and short temporal overlappings between whiskers were also included to study the precision of POm in codifying sensory patterns. Some examples are shown. Air puffs are represented by color lines. Their length reflects the duration of the signal. Temporal overlappings between sensory signals are highlighted. (B) Schematic illustration of a simulated simple tactile sequence during the exploration of an object reflecting the generation of different temporal overlappings. During the sequence, different sensory signals occur simultaneously in diverse moments generating different temporal overlappings between them. (C) Raster plots and PSTHs of representative POm responses evoked by temporal overlappings illustrated in the tactile sequence in B are shown. The appearance of a new signal during the presence of an existing signal was integrated by POm. This was reflected in precise increases in POm activity. These increases in sustained responses were only produced during the overlapping time between them (Overlapping period) but not during the non-overlapping time (Non-overlapping period of response). Color lines indicate the duration of the stimuli. The duration of the Overlapping and Non-overlapping periods is also indicated. Note that the increases of POm activity during overlapping periods were sustained along the temporal overlapping. (D) Plots comparing the spike rate of all recorded units (n = 155, depicted in different colors) during the Overlapping and Non-overlapping response periods in Control and Overlapping conditions across sensory patterns. The mean firing rate was increased in the Overlapping periods (p<0.001; Wilcoxon matched-pairs test) but not in the Non-overlapping periods where the mean magnitude of responses did not change (p = 0.43; Wilcoxon matched-pairs test). Response magnitude variation (%) between Control and Overlapping conditions in Non-Overlapping and Overlapping periods (p < 0.001, Wilcoxon matched-pairs test) is also shown.

Then, we studied these effects using sensory patterns formed by diverse temporal overlappings (Fig. 4A bottom panel). We designed these patterns to simulate possible real complex stimuli or sequences of stimuli similar to those occurring in natural circumstances, such as sequential activation of multiple whiskers with partial temporal overlapping between them, simultaneous and delayed activation of different whiskers with different durations producing sequences with diverse spatiotemporal overlappings and precise sequences of long sustained activation of several neighbouring whiskers overlapped with repetitive brief stimulations of remote whiskers. Across patterns, precise changes in POm sustained activity consistent with the spatiotemporal structure of the patterns were observed. Accurate increases in POm sustained activity were produced during the overlapping time between spatial signals across the pattern. These changes in POm activity reflected the spatiotemporal structure of the sensory pattern.

Additionally, since in natural conditions the spatiotemporal structure of sensory patterns changes dynamically across the pattern, their complexity is not homogeneous along their duration. As can be appreciated in Fig. 5A, dissimilar overlappings of spatial signals can produce sensory patterns formed by different parts with diverse complexities. To investigate the capacity of POm to represent the complexity of these different parts and to obtain a better quantification of this form of encoding, more complex sensory patterns were generated by overlapping additional spatial signals (Fig. 5B). In these stimulation protocols, we selected the most complex parts (‘Core’) of overlapping periods and divided the POm response to these parts into three subperiods: Pre, Core and Post. Precise POm activity changes were found during these different times of POm response reflecting the diverse complexity between parts along the overlapping (Fig. 5). Across patterns, POm activity was significantly increased in the Core subperiod compared to Pre and Post subperiods when extra signals were temporally overlapped in the Core subperiod of the patterns (described and quantified in Fig. 5C). This demonstrated that when additional signals were overlapped increasing the complexity of the pattern, POm codified this complexity by increasing its activity during the temporal presence of these signals. We found that POm response magnitude in all overlapping periods gradually increased as more inputs were temporally overlapped in the patterns (Fig. 5D, E). Accordingly, increasing the complexity of the pattern by increasing the number of whiskers temporally overlapped, produced an enhancement of POm response magnitude. Therefore, the complexity of spatiotemporal overlappings, represented by the number of whiskers implicated, was reflected in POm activity changes.

**Fig. 5.**
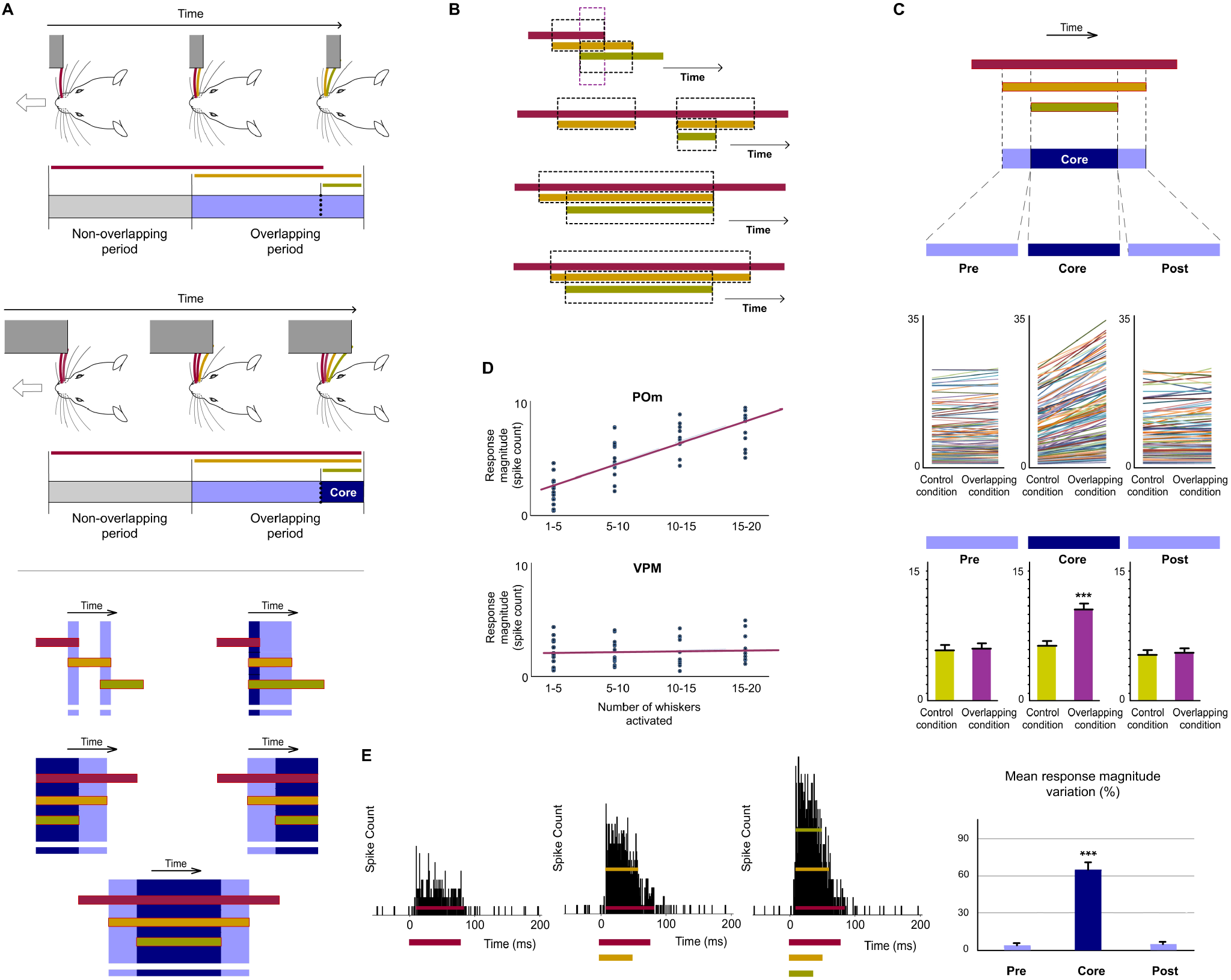
POm is highly sensitive to the dynamical spatiotemporal structure of sensory patterns and to the different complexity of their parts. (A) Multiple and constantly changing overlappings occur dynamically during natural contact with objects and surfaces during active exploration. This generates sensory patterns formed by overlappings with diverse complexities. The complexity of these overlappings determines the complexity of the patterns. As can be appreciated in these schematic illustrations of two simulated tactile sequences during the exploration of two different objects, the complexity of their corresponding Overlapping periods is different. Simple overlappings (in light blue) and more complex overlappings (in dark blue) were used to understand how POm codifies this complexity (bottom panel). The most complex parts of Overlapping periods were selected and defined as ‘Core’ parts. (B) Spatiotemporal patterns formed by more complex overlapping periods were produced by the simultaneous application of a third air puff activating additional whiskers. Different complex spatiotemporal overlappings were generated by varying the onset and duration of the third air puff and the number of extra whiskers activated by this air puff. (C) POm responses during Overlapping periods were divided in three subperiods (Pre, Core and Post). They were compared before (Control condition) and after the application of the third air puff (Overlapping condition). Data comparing the spike rate of all recorded units (n = 101, depicted in different colors) in these conditions during Pre, Core and Post subperiods across sensory patterns and the total mean response magnitude in both conditions in these subperiods are described. Mean response magnitude variation (%) in the three subperiods is also shown. POm response magnitude was significantly increased in the Core subperiod compared to Pre and Post subperiods when extra signals were temporally overlapped in the Core subperiod of the patterns (p < 0.001, One-way ANOVA). (D) Correlation between POm response magnitude and the number of whiskers simultaneously overlapped. Data from VPM are also shown for comparison. POm: Pearson correlation coefficient, r = 0.79, p < 0.001, n = 51 in 10 rats; VPM: r = 0.19, p < 0.001, n = 45 in 8 rats. Note the profound functional difference between these nuclei. (E) PSTHs for a representative example showing that POm response magnitude in Overlapping periods gradually increased as more whiskers were temporally overlapped. This shows that increasing the complexity of overlappings by increasing the number of whiskers temporally overlapped produced an enhancement of POm response magnitude. Note that the temporal structure of these different sensory events is reflected in their corresponding POm responses.

Together, these results showed that the dynamical spatiotemporal structure of sensory patterns and the different complexity of their parts were accurately reflected in precise POm activity fluctuations. This indicates that POm uses these effects to encode sensory patterns. Importantly, we observed that POm generated very similar patterns of integrated activity when different whiskers were activated by the same stimulation protocol. This finding is in agreement with the less accurate somatotopy of the nucleus and suggests that the function of POm integration is not the combined representation of specific whiskers but the encoding and extraction of generic sensory patterns from the entire vibrissal array.

#### 2.3. No sustained activity in VPM. Stimuli overlapping did not alter whisker responses in VPM

To complement the analyses described above, whisker-evoked responses in VPM were examined using stimuli with different durations (Fig. 6). Consistent with published data (Diamond et al., 1992; Simons 1995) VPM responses showed high spatial resolution (mean receptive field size: 2.3 ± 0.7 whiskers; n = 44). In contrast to POm responses and in agreement with previous findings (Castejon et al. 2016), VPM responses did not show sustained response patterns. Therefore, response modes differed drastically between these nuclei. POm was persistently activated during whisker stimulation, whereas VPM was only transiently activated at the onset of stimuli (Fig. 7A). Long stimuli usually evoked an onset response at the beginning of the stimulus and an offset response at the end but we did not find sustained responses during stimulus presence in VPM.

**Fig. 6.**
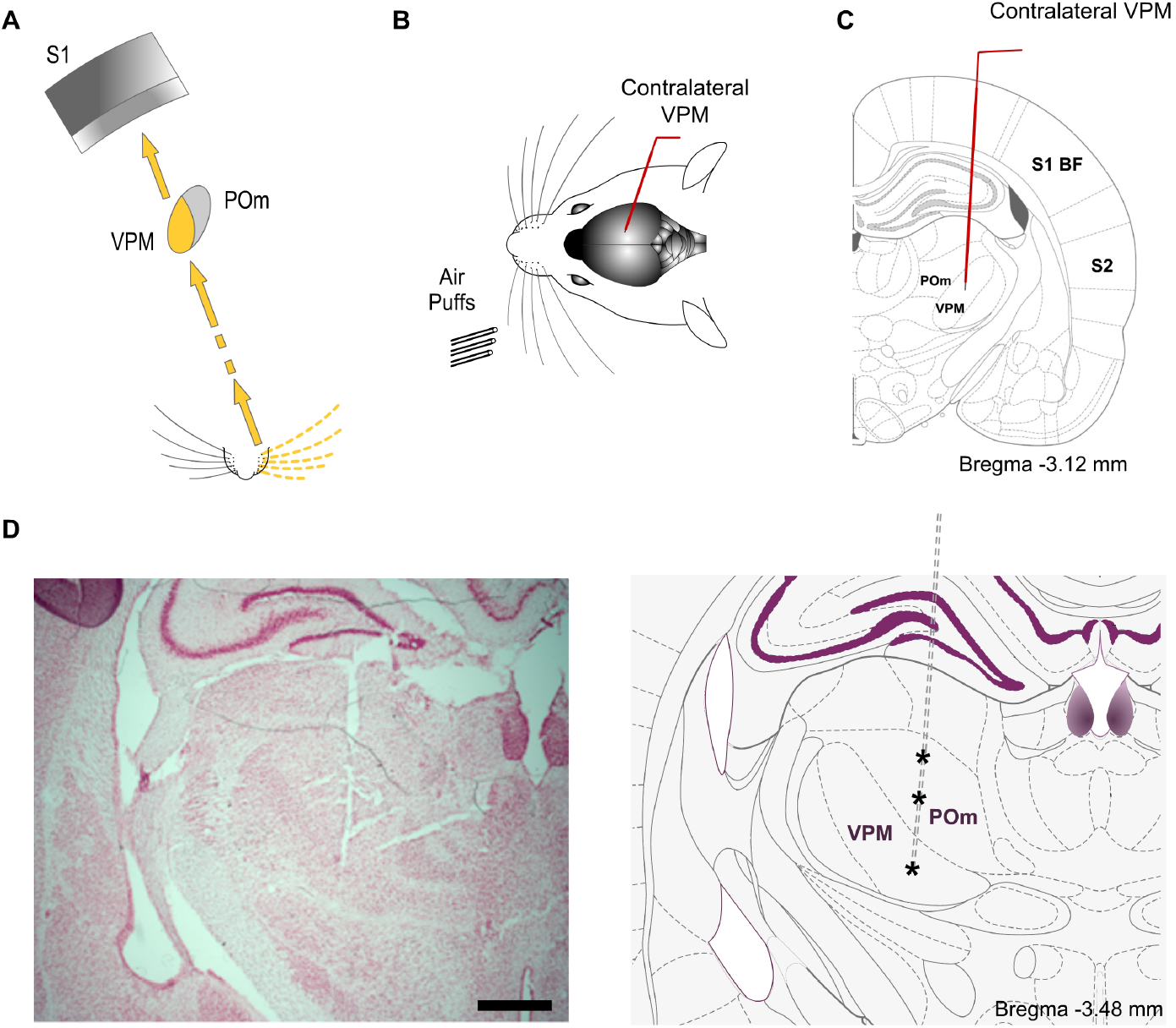
VPM. (A) Schematic illustration of the lemniscal pathway. (B) Schematic drawing displaying the sensory stimulation via patterns of multiwhisker deflections. Recordings were made in the contralateral thalamic VPM nucleus. (C). Coronal section illustrating a recording electrode inserted into the VPM. (D) The left panel shows a representative Nissl stained coronal section displaying the location of the sequence of recording sites (indicated by asterisks in the other panel) in POm and VPM and the track left by the electrode. An atlas schematic reconstruction of this coronal section is shown in the right panel (Paxinos and Watson 2007). Scale bar, 1 mm.

**Fig. 7.**
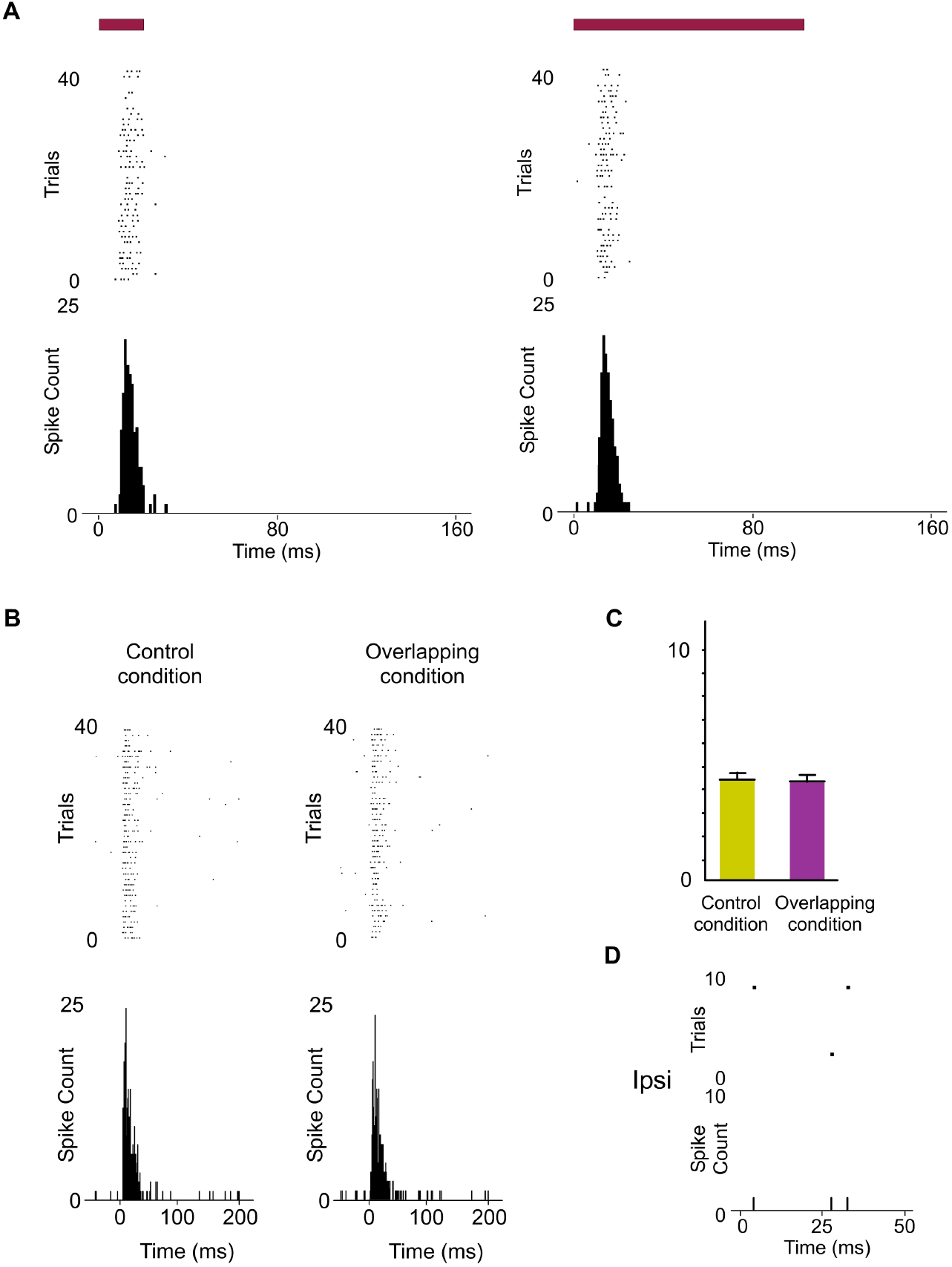
Response modes differed drastically between VPM and POm. POm sustained versus VPM transient responses. (A) VPM responses were not sustained along stimulus presence. They were transient responses just to the onset of stimuli. Raster plots and PSTHs showing multi-unit VPM transient responses evoked by different stimulus duration (20 ms and 100 ms). Note that VPM responses do not allow the discrimination between different durations of the same stimulus. Red color lines indicate the duration of the stimulus. (B) Raster plots and PSTHs showing that VPM response to simultaneous multiwhisker activation (Overlapping condition) is very similar to individual whisker activation alone (Control condition). (C) The mean response magnitude across units (n = 96) recorded in VPM did not change by the overlapping of spatial signals (−2 %, p = 0.17, Wilcoxon matched-pairs test). (D) In contrast to POm, VPM did not respond to ipsilateral stimuli (described later).

Since VPM responses were transient lasting tens of milliseconds, they seem to be excessively short for integrating over multiple whisks or longer sensory events. This suggests that temporal integration in VPM is comparatively weak. To corroborate this, we characterized VPM responses delivering spatiotemporal patterns of multiwhisker activation. Consistently across animals (n = 15), we did not find a significant change of VPM responses by multiwhisker stimuli application (Fig. 7B, see also Fig. 5E).

Together, our results show that VPM responses are different from those of POm neurons and suggest significant functional differences between POm and VPM thalamic nuclei in the processing of complex stimuli.

### 3. POm integration of spatiotemporal overlapping bilateral events

#### 3.1. POm sustained responses to ipsilateral whisker stimulation

Recently, we have described for first time that POm is also able to respond to tactile stimulation of ipsilateral whiskers (Castejon and Nunez 2020). We also showed that this nucleus is implicated in the integration of bilateral signals. Here, to study these phenomena in more detail, we recorded POm responses to ipsilateral and contralateral stimulation and confirmed that POm is able to respond to tactile stimulation of ipsilateral whiskers (Fig. 8). These experiments showed multiwhisker ipsilateral receptive fields (mean receptive field size: 9.3 ± 2.7 whiskers; n = 42) and demonstrated that POm is not only characterized by broad contralateral receptive fields but also by broad ipsilateral ones. Across recorded units (n = 90), the ipsilateral responses were weaker in magnitude than contralateral responses (Fig. 8C, D) and longer in latency (mean response onset latency: 22.31 ± 1.28 ms versus 11.36 ± 0.82 ms). This difference between ipsi- and contralateral response onset latencies (~ 10 ms, Fig. 8F) suggests that ipsilateral sensory information is mediated by different pathways.

**Fig. 8.**
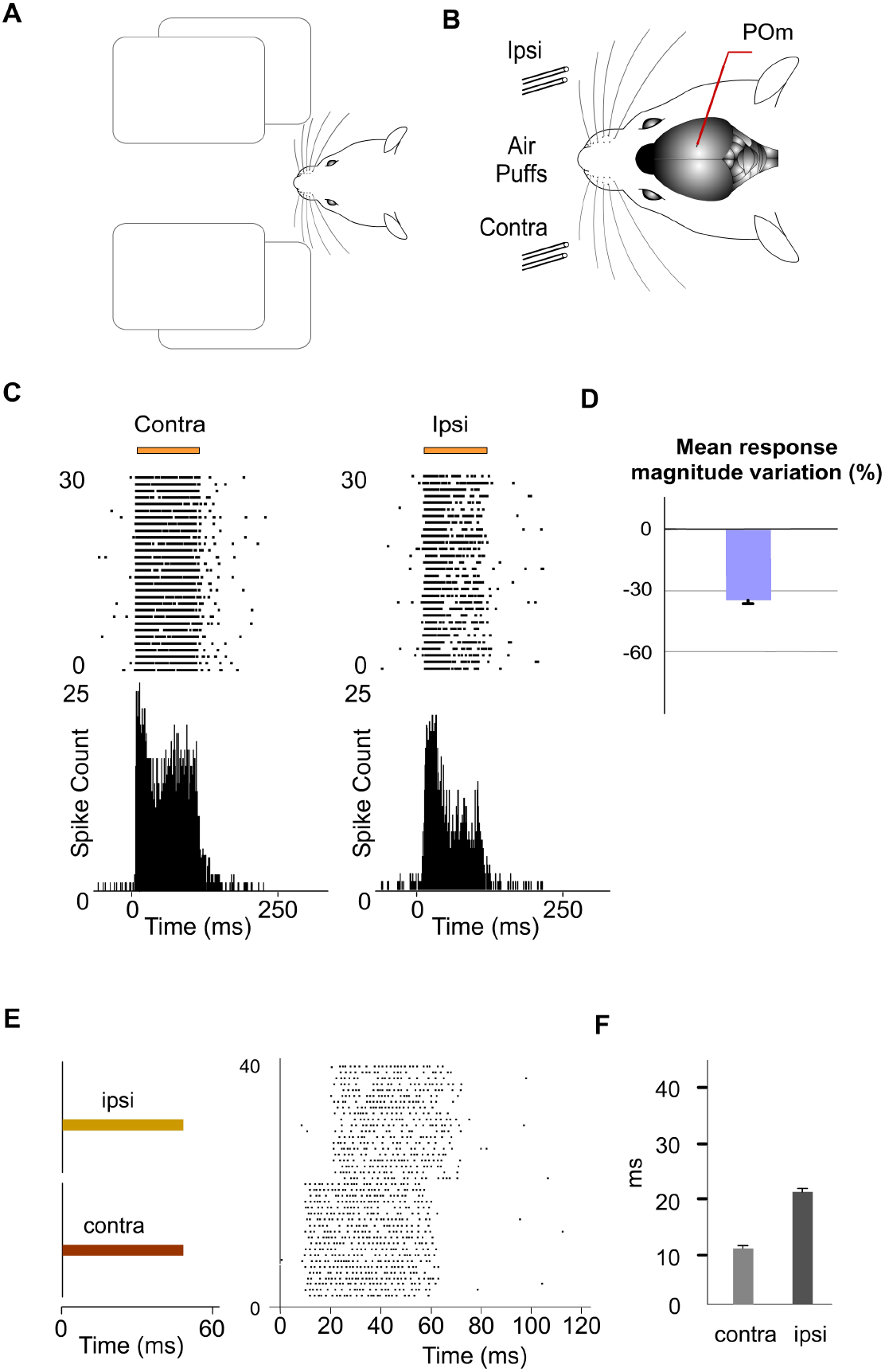
POm responses to tactile stimulation of ipsilateral whiskers. (A) Tactile information from each side of the face is needed when rodents are exploring tunnels, discerning holes and apertures or detecting their width and shape. (B) Schematic drawing displaying the sensory stimulation via patterns of contra-, ipsi- or bilateral multiwhisker deflections. Recordings were made in the POm nucleus. (C) Raster plots and PSTHs showing sustained POm responses evoked by contra- and ipsilateral multiwhisker stimulation (30 trials shown for each stimulus). Orange lines indicate the duration of the stimulus. Note that although the ipsilateral response was weaker in magnitude, the capacity to codify the duration of the stimulus remained robust. (D) Mean ipsilateral response magnitude was significantly less strong than contralateral one (p < 0.001, Wilcoxon matched-pairs test, n = 90 units). (E) Response onset latencies were longer for ipsilateral than contralateral whisker stimulation. Contralaterally evoked responses were ~10 ms faster in latency than ipsilaterally evoked ones. Color lines indicate the duration of the stimulus. Note that POm responses also lasted the duration of the ipsilateral stimulus. (F) Mean response onset latencies of contra- and ipsilateral responses are shown (n = 90 units).

Next, using air-puffs that varied in duration, we found that POm has also the capacity to sustain its activity to encode and represent tactile input duration of ipsilateral stimuli. Although the ipsilateral response was weaker in magnitude, the capacity to codify the duration of the stimulus remained robust as reported in the raster plots of a representative response in Fig 8C, E. The duration of the ipsilateral stimuli did not alter the mean onset latency of responses (One-way ANOVA, p = 0.79). These findings were consistent across all animals (n = 10).

#### 3.2. POm integration of spatiotemporal overlapping ipsilateral events

Next, we investigated the implication of POm in the encoding of complex ipsilateral stimuli or patterns of stimuli. Accordingly, we studied POm responses to ipsilateral multiwhisker activation protocols producing different sensory patterns of spatiotemporal overlappings. Across these patterns, the firing rate was significantly increased in the Overlapping periods (Fig. 9). These increases were also sustained along the temporal overlappings (Fig. 9D). However, in the Non-overlapping periods, the magnitude of responses did not change. Again, we found precise changes in POm activity consistent with the spatiotemporal structure of the ipsilateral patterns (Fig. 9 D). These findings were consistent across all animals (n = 11).

**Fig. 9.**
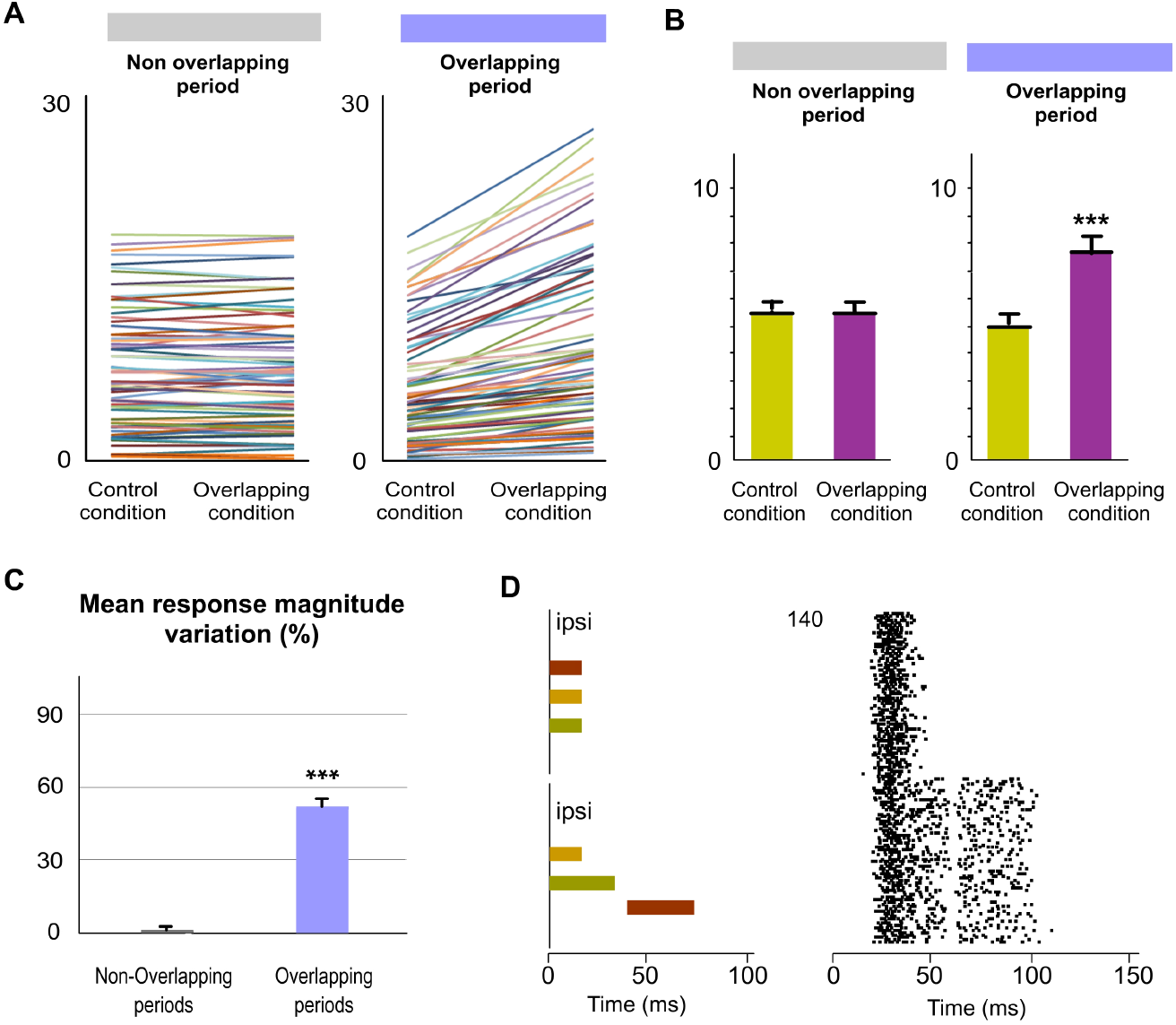
POm integration of ipsilateral overlapping events. (A) Data showing the quantification of the facilitative integration during overlappings of ipsilaterally evoked signals. Plots comparing the spike rate of all recorded units (n = 80, depicted in different colors) during the Overlapping and Non-overlapping response periods in Control and Overlapping conditions across sensory patterns. (B) The mean firing rate was significantly increased in the Overlapping periods (p<0.001; Wilcoxon matched-pairs test) but not in the Non-overlapping periods where the mean magnitude of responses did not change (p = 0.97; Wilcoxon matched-pairs test). (C) Mean response magnitude variation (%) between Control and Overlapping conditions in Non-Overlapping and Overlapping periods (p < 0.001, Wilcoxon matched-pairs test). (D) Peri-stimulus time raster plots of representative POm responses to two different overlapping protocols of multiwhisker ipsilateral stimulation (70 trials shown for each protocol). An increase in POm response magnitude can be appreciated as more inputs were temporally overlapped in these protocols. Note that the general temporal structure of the sensory pattern, the temporal position of its components (depicted in different colors) and their corresponding overlappings are reflected in the structure of POm response.

We also characterized VPM thalamic responses delivering the same spatiotemporal patterns of ipsilateral multiwhisker activation in 7 rats. In contrast to POm, VPM did not respond to ipsilateral stimuli or to ipsilateral overlapping protocols (Fig. 7D). Since the integration of tactile information from the two sides of the body is fundamental in bilateral perception, our results suggest a different implication of these thalamic nuclei in this function.

#### 3.3. Transmission of integrated sensory activity between both POm nuclei

These findings raise the question of by which route(s) is the ipsilateral information relayed to POm. It is known that POm is subcortically innervated by the Principal (Pr5) and Interpolar nuclei (SpVi) but we did not find evoked responses to contralateral whisker stimulation in these trigeminal nuclei (11 rats), therefore POm responses to ipsilateral whiskers stimulation were not driven by ascending peripheral activity conveyed directly via the trigeminal complex. In agreement with this finding, the difference between latencies in POm to contralateral and ipsilateral stimuli suggests that ipsilateral information is not received by POm directly from the periphery. The delay (10 ms) that we observed for ipsilateral information suggests the indirect transfer of ipsilateral information between hemispheres from the other POm. To test this possibility, unilateral overlapping sensory patterns were applied while extracellular recordings were performed in both POm nuclei simultaneously in 6 rats (Fig. 10). We found that responses evoked by these unilaterally applied overlapping sensory patterns were similar in both nuclei. In agreement with our data described above, they showed different onset latencies (Fig. 10B) suggesting that evoked activity first arrives at the contralateral POm and is then transferred to the ipsilateral POm in the other hemisphere. This would indicate that both POm nuclei are mutually connected forming a POm–POm loop.

**Fig. 10.**
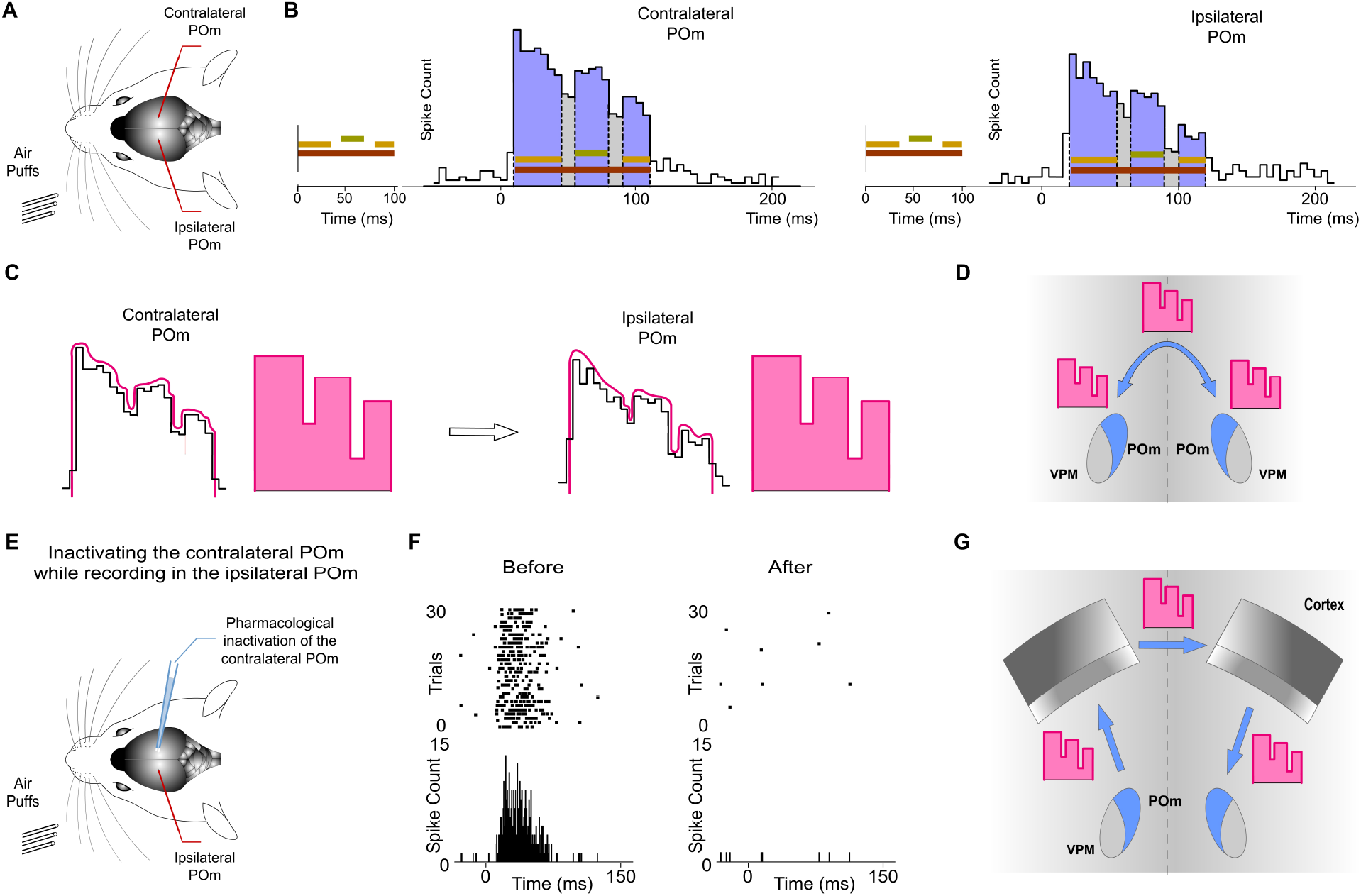
Ipsilateral sensory information is received by POm interhemispherically from the other POm. This activity was transmitted preserving its integrated structure across the POm-POm loop. (A) Unilateral overlapping sensory patterns were applied while extracellular recordings were performed in both POm nuclei simultaneously. (B) POm responses evoked by these unilaterally applied patterns showed different onset latencies but similar integrated structure in the contra- and ipsilateral POm nuclei. As can be appreciated in these PSTHs (bin width 5 ms) of POm responses, precise changes in POm activity caused by the integration of overlapping signals were similar in the contra- and ipsilateral POm nuclei. Note that the onset latency was longer in the ipsilateral POm. (C) PSTHs in B show that precise changes (fluctuations represented by lines in pink) in POm sustained activity caused by the integration of overlapping stimuli were precisely conserved. Therefore, these patterns of integrated information encoded by POm were interhemispherically transmitted to the other POm preserving their integrated structure. These fluctuations of POm activity can be considered as ‘structured patterns of integrated activity’ (represented by schematic patterns in pink). (D) These ‘structured patterns’ are transmitted across the POm-POm loop. (E) POm responses to ipsilateral stimulation were studied before and after pharmacological deactivation by muscimol (1 mg/ml) injection in the opposite POm. (F) POm responses to ipsilateral stimulation were abolished when the opposite POm was pharmacologically deactivated. (G) The latencies suggest that the POm-POm loop could be formed by a thalamocortical-callosal-corticothalamic route.

Importantly, precise changes in POm activity caused by the integration of overlapping stimuli reflecting the spatiotemporal structure of the sensory pattern were precisely conserved (Fig. 10B). Therefore, these patterns of integrated information encoded by one POm were transmitted through the loop to the other POm preserving their integrated structure (Fig. 10C).

To confirm that ipsilateral activity reaches one POm from the other POm, we pharmacologically deactivated one of them by muscimol (1 mg/ml) injection (Fig. 10E). We found that evoked responses in the second POm to ipsilateral stimulation were abolished in the majority of cases (4 out of 6 rats; Fig. 10F). However, they were almost abolished but not completely eliminated in 2 cases (−91 %, p < 0.001). This residual activity may be attributed to an incomplete deactivation of the opposite POm or a transmission of ipsilateral activity by an alternative pathway (i.e., collicular commissure). However, we observed that the sustained patterns of integrated activity evoked by ipsilateral sensory patterns were abolished in all animals. This indicated that these sustained patterns of integrated information were received from the other POm. Together, these findings demonstrated a transmission of integrated sensory activity between both POm nuclei through a functional POm-POm loop.

#### 3.4. POm integration of spatiotemporal overlapping bilateral events

Finally, since bilateral sensory events occur concurrently producing different overlappings, the next question to investigate was whether POm would be able to codify these complex bilateral dynamics. It would require the precise integration of contralateral sensory inputs from the brainstem and ipsilateral sensory inputs from the other POm. To investigate the implication of POm in these intricate computations, we applied spatiotemporal overlapping patterns of bilateral multiwhisker stimulation simulating possible natural bilateral sensory events (Fig. 11A). We measured the responses of the nucleus to the overlapping patterns and found that POm precisely integrates tactile events from both sides. We found that precise changes in the spatiotemporal structure of bilateral events evoked different patterns of POm integrated activity. Across sensory patterns, the firing rate was increased in the Overlapping periods (quantified in Fig. 11) but not during the non-overlapping time. This indicates that the time shared by overlapping ipsi- and contralateral stimuli is encoded by POm activity. These increases in firing rate were sustained along the Overlapping periods (Fig. 11B). Similar effects were found stimulating identical (mirror) or different whiskers (nonmirror whiskers) on both sides. This is in agreement with the less accurate somatotopy of the nucleus and again indicates that the function of POm integration is not the combined representation of specific whiskers but the generic encoding of sensory patterns integrated from both whisker pads. Moreover, during Overlapping periods, facilitation of responses was found as more ipsilateral, contralateral or bilateral temporal overlapping inputs were added showing that the complexity of the bilateral spatiotemporal overlappings is replicated in POm activity variations. These findings were consistently found across animals (n = 17).

**Fig. 11.**
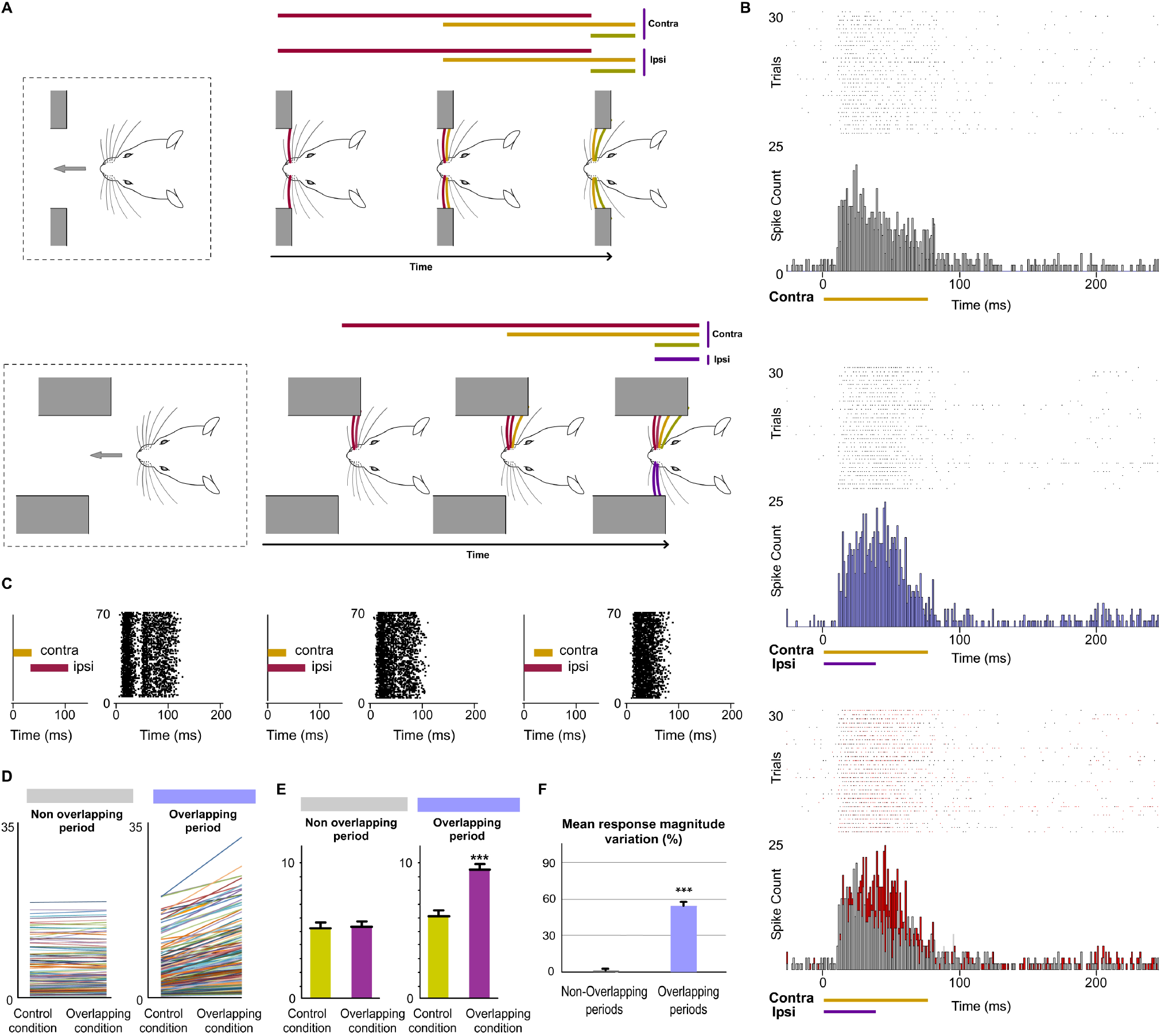
POm integration of bilateral events. (A) As can be appreciated in these two simulated exploratory tactile sequences, many whiskers on both sides are sequentially deflected simultaneously. In our experiments, bilateral sensory patterns were generated by the application of contralateral and ipsilateral stimuli activating different subsets of whiskers on both sides of the face at mirror (matched whiskers on each side) and nonmirror positions. (B) Raster plots and PSTHs showing demonstrative POm responses evoked by contralaterally (top) and bilaterally (middle) evoked overlapping stimulation using stimuli with different durations (color lines indicate the duration of the stimuli). Note that the time shared by these bilateral overlapping inputs is encoded by a precise increase in POm activity. This increment is depicted in red in the bottom panel. (C) The time interval between contralateral and ipsilateral stimuli was varied to produce different overlappings and to study their integration. Raster plots of representative POm responses to different bilateral overlapping stimulation protocols are shown. Note that since the onset latency of ipsilateral responses is ~10 ms longer than contralateral responses, the temporal interval between contralateral and ipsilateral signals determines their integration and the structure of the response. As can be appreciated in the left panel, no integration was observed when the contra- and ipsilateral stimuli were not temporally overlapped. (D) Data showing the quantification of the facilitative integration during overlappings of contralaterally and ipsilaterally evoked signals. Plots comparing the spike rate of all recorded units (n = 187, depicted in different colors) during the Overlapping and Non-overlapping response periods in Control and Overlapping conditions across bilateral sensory patterns. (E) The mean firing rate was significantly increased in the Overlapping periods (p<0.001; Wilcoxon matched-pairs test) but not in the Non-overlapping periods where the mean magnitude of responses did not change (p = 0.41; Wilcoxon matched-pairs test). (F) Mean response magnitude variation (%) between control and overlapping conditions in Non-Overlapping and Overlapping periods (p < 0.001, Wilcoxon matched-pairs test).

Crucially, we found that the temporal interval between bilateral stimuli is critical to bilateral integration of sensory information. The delay (~10 ms) that we observed for ipsilateral information determines the interaction between bilateral stimuli (Fig. 11C).

#### 3.5. POm nuclei are mutually connected through the cortex

Our results prompt the question of by which route(s) is sensory information transferred from one POm to the other. Cortical responses to ipsilateral whisker stimulation have been described in the somatosensory cortex (Shuler et al. 2001; Debowska et al., 2011). Therefore, ipsilateral activity seems to arrive at the contralateral POm by crossing the corpus callosum and descending from the cortex. Since POm receives strong innervation from corticofugal projection neurons in S1 (Hoogland et al., 1991; Bourassa et al., 1995; Veinante et al., 2000b), it is then possible that ipsilateral sensory stimulation could produce the activation of these descending corticofugal projections. This could have important implications on the integration of cortical inputs by POm and suggests that ipsilateral stimulation can be used to study the nature and content of the messages travelling through these corticofugal projections. Moreover, it has been proposed that functioning of higher-order nuclei, including POm, are determined by these descending corticofugal projections. To confirm that ipsilateral activity reaches POm via corticothalamic axons and to investigate whether these thalamic capacities are generated or mediated by cortical influence, we studied POm response properties before and after pharmacological deactivation of S1 (of the same hemisphere) by lidocaine (10%) or muscimol (1 mg/ml) application. As a control, we simultaneously recorded neuronal activity in the injected area. The electrodes were placed in the infragranular layer and the inactivation was confirmed by the absence of spontaneous and evoked activity. We found that POm responses to ipsilateral stimulation were almost abolished when S1 was inactivated (−89 %, p < 0.001, paired t-test, n = 8 rats; Fig. 12B). However, since the attempt to pharmacologically inactivate S1 can produce its partial deactivation and also can affect surrounding cortical areas, we confirmed our result by cortical lesion. The lesion was restricted to S1 and included superficial and deep layers of this area. This approach showed similar results. POm responses to ipsilateral stimulation were almost eliminated (−92. p < 0.001, paired t-test, n = 6 rats).

**Fig. 12.**
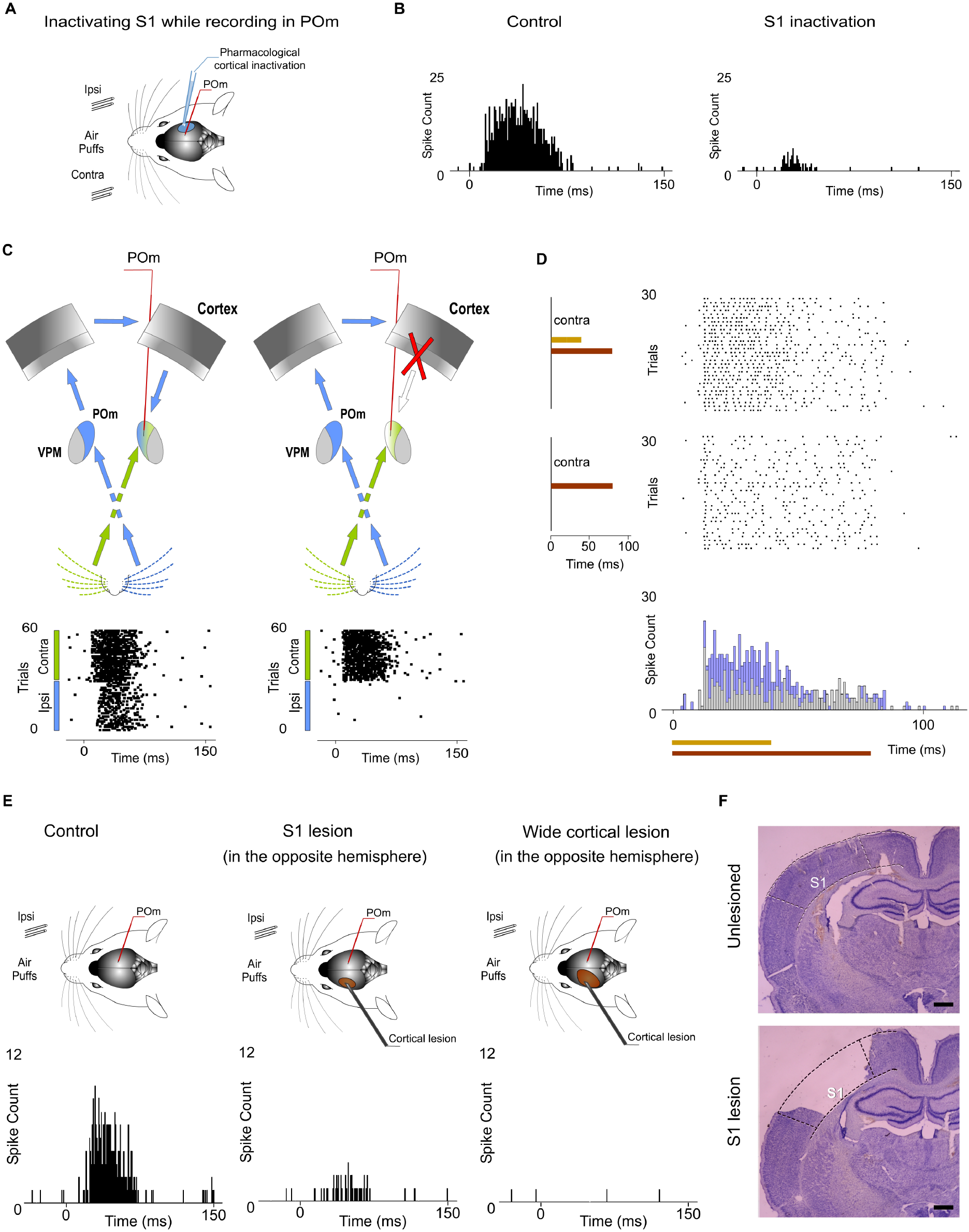
The POm-POm loop. Ipsilateral and contralateral POm nuclei are mutually connected through the cortex. (A) POm responses were recorded before and after S1 inactivation. (B) POm responses to ipsilateral whisker stimulation were almost abolished when S1 was inactivated or lesioned. PSTHs of representative POm responses before and after S1 inactivation. (C) Schematic illustration representing the transmission of contralateral and ipsilateral information to POm. As can be appreciated in the corresponding raster plots, POm response to ipsilateral whisker stimulation was eliminated when the POm-POm loop was interrupted by cortical removal. Note that POm response to contralateral stimulation was still present in this condition. (D) Cortical removal did not affect the capacity of POm to sustain its activity to codify stimuli duration and its capacity to integrate overlapping stimuli as can be appreciated in these raster plots and PSTHs of POm responses recorded during this condition. (E) PSTHs of representative POm responses in control, S1 lesion (in the opposite hemisphere) and wide cortical lesion (in the opposite hemisphere) conditions are shown. An almost complete reduction but not a total elimination of ipsilateral responses was produced by the S1 lesion. POm responses to ipsilateral stimulation were completely eliminated only when a wider cortical extension was lesioned. (F) Example histology showing the removal of S1 from a lesioned animal. An unlesioned example is also shown for comparison. Scale bars, 1 mm.

POm also receives cortical projections from M1 and S2 (Alloway et al. 2008; Liao et al. 2010). However, we did not find a significant reduction of ipsilateral responses when only M1 and S2 were lesioned (−7 %, p = 0.32, paired t-test, n = 4 rats). Together, these findings are in agreement with previous studies showing that cortical ‘driver’ input (Sherman and Guillery 1998) to POm originates almost exclusively from S1 (Veinante et al. 2000b).

When a wider cortical extension was lesioned including S1, M1 and S2, POm responses to ipsilateral stimulation was completely abolished in the majority of cases (4 out of 6 rats). However, we still found very small responses to ipsilateral stimulation in 2 cases (−94 %, p < 0.001). Importantly, we found robust contralateral whisker-evoked responses in POm even when the cortex of the same hemisphere was inactivated, which indicates a direct ascending input from the periphery. Cortical inactivation slightly affected the magnitude of POm responses to contralateral whiskers. We only found a minimal reduction of spikes (−8 %; paired t-test, p = 0.06, n = 6 rats; Fig. 12C). Furthermore, using air-puffs that varied in duration to stimulate contralateral whiskers, we found that cortical inactivation did not affect the capacity of POm to sustain its activity to codify stimuli duration Fig. 12D). We also measured POm responses to contralateral overlapping sensory patterns in this condition and found that cortical inactivation did not affect the capacity of POm to integrate overlapping stimuli (Fig. 12D). On the basis of these results, we conclude that POm does not inherit these capacities from cortical influence.

Together, these results demonstrated that ipsilateral and contralateral POm nuclei are mutually connected through the cortex by showing that ipsilateral activity reaches POm via descending parallel corticofugal projections mainly from S1 of the same hemisphere. But, by which route(s) is the ipsilateral information relayed to S1? Since projections from POm to the cortex in the other hemisphere have not been described and since it is known that corticocortical transmission between hemispheres via callosal projections is the main route for ipsilateral sensory inputs (Shuler et al. 2001; Petreanu et al., 2007), it seems that the POm-POm loop could be formed by a thalamocortical-callosal-corticothalamic route. To test this, we studied POm responses to ipsilateral stimulation before and after deactivation of S1 in the other hemisphere by lidocaine (10%) or muscimol (1 mg/ml) application. We found an almost complete reduction but not a total elimination of ipsilateral responses (−86 %, p < 0.001, paired t-test, n = 6 rats). This was confirmed by S1 lesion (p < 0.001, paired t-test, n = 5 rats; Fig. 12E). Again, when a wider cortical extension was lesioned including S1, M1 and S2, POm responses to ipsilateral stimulation were completely abolished in the majority of cases (4 out of 5 rats). We still found residual responses to ipsilateral stimulation in one case. This remaining activity could be attributable to other cortical areas projecting to POm or to subcortical interhemispheric pathways such as the collicular commissure.

Finally, to investigate the implication of cortical layers in the processing and transmission of sustained activity between the thalamus and the cortex in the POm-POm loop, evoked responses across recorded multi-units in supra, granular and infragranular layers of S1 were examined (n = 117, n = 88 and n = 102 units respectively) using ipsi- and contralateral stimuli with different durations in 19 rats. Examination of the laminar profile of evoked activity across layers showed profound differences between them. We only found sustained responses lasting the duration of the stimulus in the infragranular layer (Fig. 13). Similar to VPM responses, supra- and granular responses were only transiently activated at the onset of stimuli (Fig. 13D). Long stimuli usually evoked an onset response at the beginning of the stimulus and an offset response at the end but we did not find sustained responses during stimulus presence in these layers. These findings demonstrated different laminar profile of cortical responses. Moreover, using ipsilateral stimuli with different durations to examine the implication of these layers in the interhemispheric transfer of sustained activity, we found that the infragranular layer showed evoked sustained responses to ipsilateral stimulation. These results demonstrated different laminar implication in the processing of sustained activity and its transmission between hemispheres.

**Fig. 13.**
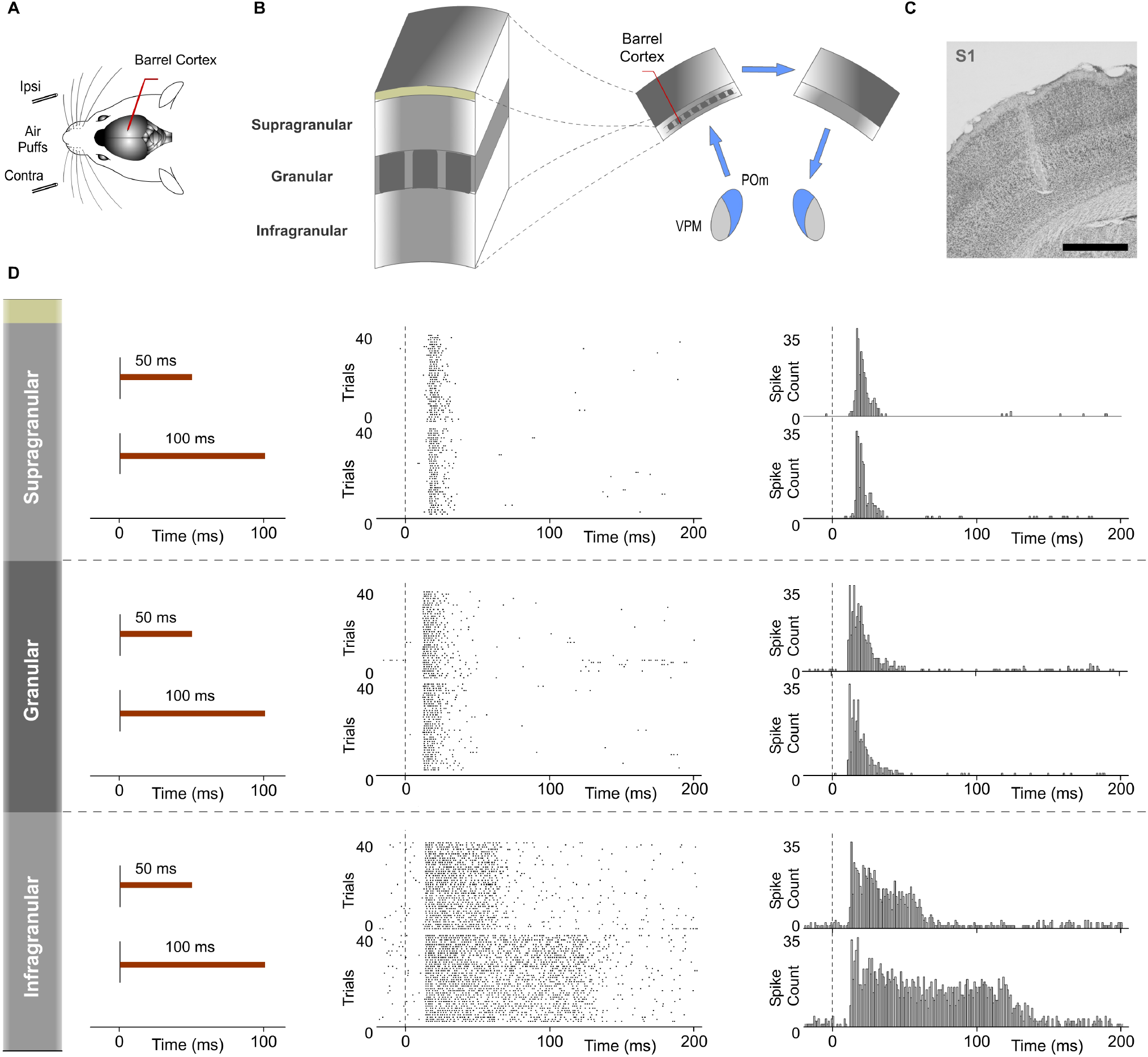
Response modes differed between cortical layers in S1. Sustained responses along stimulus presence were found in the infragranular layer but not in granular and supragranular layers. (A) Recordings were made in the barrel cortex in S1 using ipsi- and contralateral stimuli with different durations. (B) Evoked responses in supra-, granular and infragranular layers of the barrel cortex were examined to study the laminar implication in the processing of sustained activity and its transmission across the POm-POm loop. (C) Histological section displaying the location of the sequence of recording sites across cortical layers in S1 and the track left by the electrode. Scale bar, 1 mm. (D) Raster plots and PSTHs showing typical responses in supragranular, granular and infragranular layers evoked by contralateral stimuli with different durations (50 ms and 100 ms). Note that evoked responses in granular and supragranular layers were transient just to the onset of stimuli and that they do not allow the discrimination between different durations of the same stimulus. Color lines indicate the duration of the stimulus.

## Discussion

### 1. The role of higher-order sensory thalamus in the encoding of complex sensory patterns

The function of the higher-order sensory thalamus remains unclear. Here, we propose the hypothesis that an important function of higher-order thalamic nuclei could be the encoding of complex sensory patterns. Their functional implication may allow the processing and extraction of patterns and regularities from the sensory input. They could allow sensory systems to generate a representation of these dynamics. Our findings describing the different implication of POm and VPM in the processing of complex stimuli are in agreement with this proposal. This also needs to be confirmed in other sensory modalities.

### 2. The capacities of POm to sustain and integrate activity

How does the brain discriminate between different durations of identical stimuli? Here, we confirm previous findings (Castejon et al. 2016) by demonstrating that POm has the capacity to sustain its activity to represent tactile event duration with high accuracy. Extracting this temporal information from the sensory input is essential for the optimal extraction of information. In addition, our results show that POm is functionally implicated in the computation of overlapping spatiotemporal signals. This computation was performed by a precise facilitative integration of these signals.

Our findings show that the capacities of POm to sustain and integrate activity allow the representation of complex tactile events. Moreover, these POm capacities were still present during cortical inactivation. Accordingly, we conclude that they are not inherited from cortical influence. Importantly, these functional POm capacities are obtained from population activity (Fig. 2B). The presence of these functional capacities could be a consistent feature of higher-order thalamic nuclei.

### 3. Functional significance of POm capacities

#### POm activity fluctuations to codify patterns

Varying the spatiotemporal structure of sensory patterns, we found that POm is highly sensitive to multiwhisker activation involving complex spatiotemporal interactions. Our results show that the dynamical spatiotemporal structure of sensory patterns and the different complexity of their parts were accurately reflected in precise POm activity changes. Therefore, these precise fluctuations of POm integrated activity generate representations of these dynamics. This finding prompts a fundamental question: What could be the function of these precise thalamic activity fluctuations? Since they are composed by integrated activity generated by POm reflexing the spatiotemporal structure of the sensory input, POm integration may provide a mechanism for detecting spatiotemporal landmarks in the continuous flow of incoming sensory signals. It could be used to precisely decode sequence boundaries allowing for extraction of regularities and patterns from the flow of raw sensory information. Moreover, these fluctuations can also serve as relevant cues in sensorimotor adjustment, pattern recognition, perceptual discrimination and decision-making.

From this functional perspective, active whisking and palpation movements can intentionally optimize the number, frequency and variation of overlappings to maximize the extraction of information (i.e., regularities) from objects, surfaces and textures during their exploration. It is known that rodents use different whisking strategies changing from large-amplitude whisker movements during wide exploration to small-amplitude whisker movements at higher frequencies to increase the resolution in the extraction of detailed information. In agreement with this, we found that POm has the ability to integrate overlapping inputs with high precision even when very short duration stimuli were used (20 ms). This finding is compatible with the range of frequencies described in these animals: between 4-12 Hz (large whisker movements) and 12-25 Hz (small movements; Carvell, and Simons 1990). Accordingly, adapting the active generation of precise overlappings of specific subsets of whiskers and their frequency would allow these animals to obtain the optimal resolution necessary to solve different perceptual or tasks requirements.

#### POm is an encoder of patterns

Rodents and other mammals have the ability to detect complex patterns embedded in a continuous stream of sensory activity. Across protocols, we found that varying the spatiotemporal structure of the sensory input produced different patterns of POm integrated activity. We observed that POm generates very similar patterns of activity when different whiskers were activated by the same stimulation protocol. This finding is in agreement with the less accurate somatotopy of this nucleus and suggests that the function of POm integration is not the combined representation of specific whiskers but the encoding of generic sensory spatiotemporal patterns from the array of whiskers. Accordingly, our findings suggest that POm is a general encoder of patterns.

### 4. POm mediates bilateral sensory processing

#### POm integration of bilateral events

Although it is well described that POm encodes stimulations of the contralateral whisker pad, our results show that POm is also able to respond to tactile stimulation of ipsilateral whiskers. This finding challenges the notion that the somatosensory thalamus simply computes contralateral whisker stimuli. In addiction, we found that POm integrates signals from both whisker pads and described how this integration is generated. Importantly, we found that precise changes in the spatiotemporal structure of bilateral events evoked different patterns of POm integrated activity.

These findings can explain why the presence of tactile input from one side affects tactile processing of the other side (for example, in sensory interference paradigms) and demonstrate the implication of the higher-order sensory thalamus in the encoding of bilateral events.

#### The POm-POm loop: POm nuclei are mutually connected through the cortex

Our results show that ipsilateral activity reaches one POm from the other POm (Fig. 10). Moreover, these findings demonstrate a transmission of integrated activity between both nuclei through a functional POm-POm loop formed by thalamocortical, interhemispheric and corticothalamic projections. We confirmed this interhemispheric pathway by inactivating different areas of the cortex in both hemispheres (Fig. 12) and demonstrating that ipsilateral activity mainly reaches POm via S1 but not exclusively. Therefore, POm nuclei are mutually connected forming a complex network of parallel thalamocortical, interhemispheric and corticothalamic projections (Fig. 14A). In agreement with this finding, it is anatomically well described that MI and S2 also receives thalamocortical projections from POm (Ohno et al. 2012), that they are respectively interhemispherically connected (Carvell and Simons, 1987, Kinnischtzke et al. 2014) and that they have corticothalamic projections to POm (Alloway et al. 2008; Liao et al. 2010). Moreover, POm is a strong driver of activity in S2 and M1 (Theyel et al., 2010; Castejon et al. 2016; Casas-Torremocha et al. 2019) and bilateral sensory responses in S2 have also been described (Debowska et al., 2011). However, it is important to note that the remaining ipsilateral activity that we found after S1 deactivation did not allow for the encoding and represention of ipsilateral tactile event duration or its spatiotemporal structure with high accuracy. This indicates that although different cortical areas are parallelly implicated in the POm-POm loop, the robustness of this encoding is supported by S1. Furthermore, since S1 receives intra- and interhemispheric projections from these cortical areas (Porter and White 1983; Carvell and Simons, 1987; Kinnischtzke et al. 2014), it is also possible that S1 could collect activity from different cortical regions (mostly from the other S1) and funnel this information via corticothalamic projections to POm of the same hemisphere (Fig. 14A). Consistent with this idea, it is known that motor and S2 cortical regions elicited strong direct input to L6b and L5 in S1 (Mao et al. 2011; Zolnik et al., 2020). Moreover, corticofugal projections from L6b to POm have also been confirmed (Bourassa et al. 1995; Hoerder-Suabedissen et al. 2018). L6b receives substantial innervation from the contralateral sensorimotor cortical areas producing a contralateral drive to this layer (Zolnik et al. 2020). Here, we found that VPM did not respond to ipsilateral stimuli. Unlike POm, VPM does not receive cortical input from L5 or L6b corticofugal projections in S1 (Hoogland et al. 1991; Hoerder-Suabedissen et al. 2018).

**Fig. 14.**
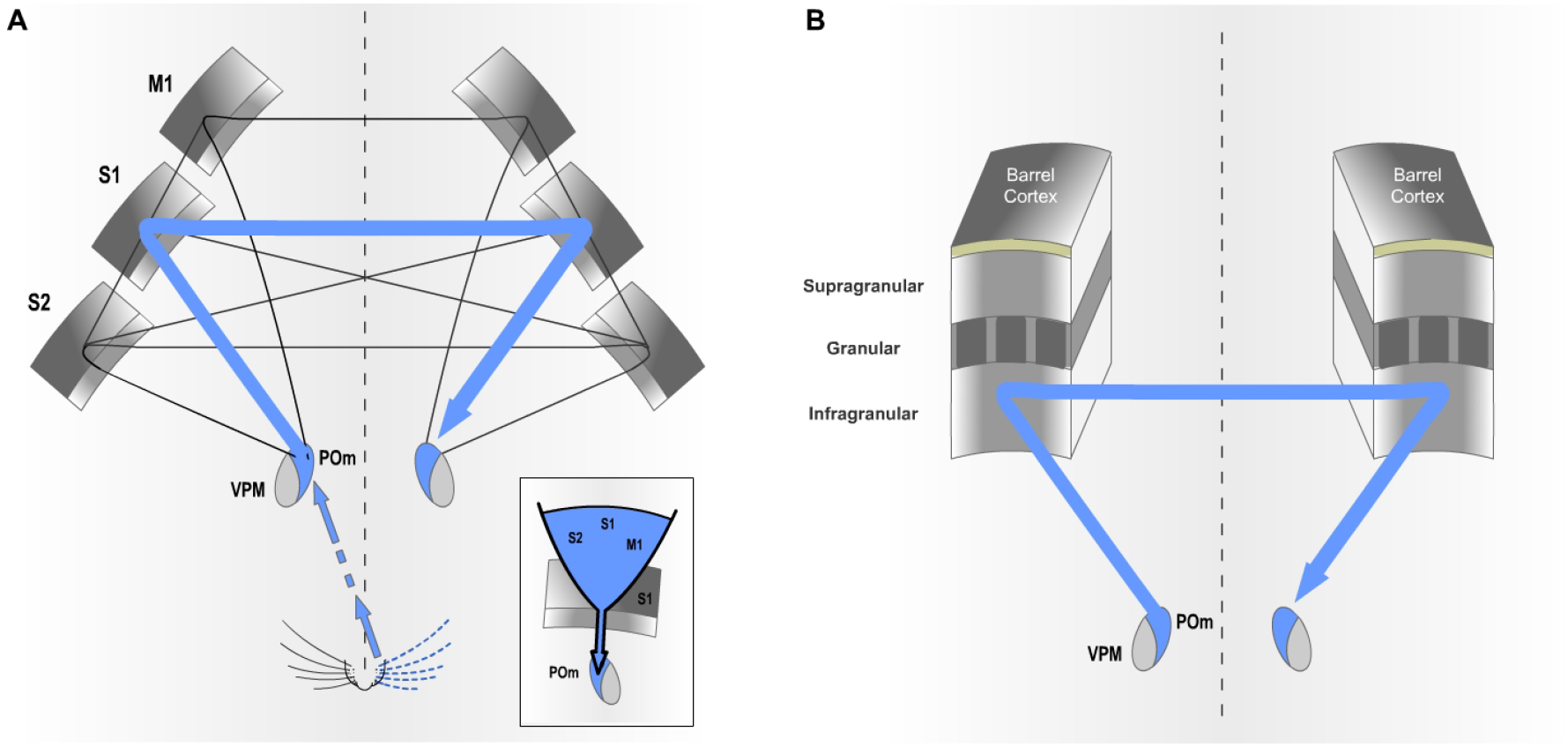
The POm-POm loop is formed by a functional network of parallel thalamocortical, interhemispheric and corticothalamic projections. (A) Although different cortical areas are implicated, our results revealed that S1 plays a protagonistic role in this functional loop. The inset illustrates the idea that S1 may act as a funnel collecting activity from different areas and sending this information via corticothalamic projections to POm of the same hemisphere. This complex interhemispheric loop allows bilateral integration in the thalamus and is implicated in the bidirectional transmission of sustained activity between the higher-order thalamus and the cortex. This transmission of sustained activity can be functionally implicated in cognitive processes. (B) Our results indicate that the transmission of sustained activity across the POm-POm loop is supported by the infragranular layer. This loop could be also present in other sensory modalities.

Importantly, our results also demonstrated different laminar implications in the processing of sustained activity and its transmission between hemispheres (Fig. 13). We only found sustained responses lasting the duration of the stimulus in the infragranular layer. Similar to VPM responses, supra- and granular responses were only transiently activated at the onset of stimuli but we did not find sustained responses along stimulus presence. Using ipsilateral stimuli with different durations to examine the implication of these layers in the interhemispheric transfer of sustained activity, we found that only the infragranular layer showed evoked sustained responses to ipsilateral stimulation. These findings demonstrate that the transmission of sustained activity across the POm-POm loop is supported by the infragranular layer (Fig. 14B).

In addition, we found that POm constantly integrates bilateral sensory information and that ipsilateral activity reaches POm via corticofugal projections mostly from S1. This in agreement with previous findings showing the convergence of ascending driver inputs from the periphery and descending driver inputs from L5 of S1 in POm (Groh et al. 2014; Castejon et al. 2016). We found that the temporal interval between contra- and ipsilateral stimuli was critical to bilateral integration of sensory information. The delay (~10 ms) that we observed for ipsilateral information determined the interaction between bilateral stimuli (Fig. 11C). Also consistent with our findings is that the activation of corticofugal projections from L5 in S1 by optogenetic stimulation increases ascending sensory responses within a well-defined time window (Groh et al. 2014). In our study, we took advantage of the fact that ipsilateral sensory stimulation produced the activation of these corticothalamic fibers to investigate, in more physiological conditions, the functional interaction of these two streams in the integration of bilateral events and the implication of these corticothalamic projections in POm functioning.

### 5. Structured patterns of integrated activity as ‘Templates’ for cognitive functions

Our results raise additional questions of interest with regard to the possible functional implications of the sustained thalamic patterns of integrated activity. We found that the sequences of precise fluctuations of POm activity constitute ‘structured patterns of integrated activity’. We suggest that the patterns of structured activity generated by POm may be functional building blocks of perception and information extraction. Accordingly, they may serve the function of transforming raw sensory input into useful information by allowing the extraction of relevant patterns from the continuous sensory flow (Fig 15). In agreement with this idea, rodents have the capacity to locate, identify and discriminate shapes, objects and other sensory patterns in their environment with very high precision. But what functional roles could these integrated patterns play apart from perception? We propose that these patterns of structured activity may allow the conversion of sensory information into functional representations relevant to current or future cognitive tasks. Therefore, they could be selected, acquired and used as ‘Templates’ for cognitive functions. From this perspective, one important function of higher-order sensory thalamic nuclei may be the generation of perceptual units extracted from the sensory flow and converted into functional units which could be used as Templates for current or future cognitive requirements. This functional proposal provides a novel theoretical model to understand the implication of the thalamus in perception and cognition.

**Fig. 15.**
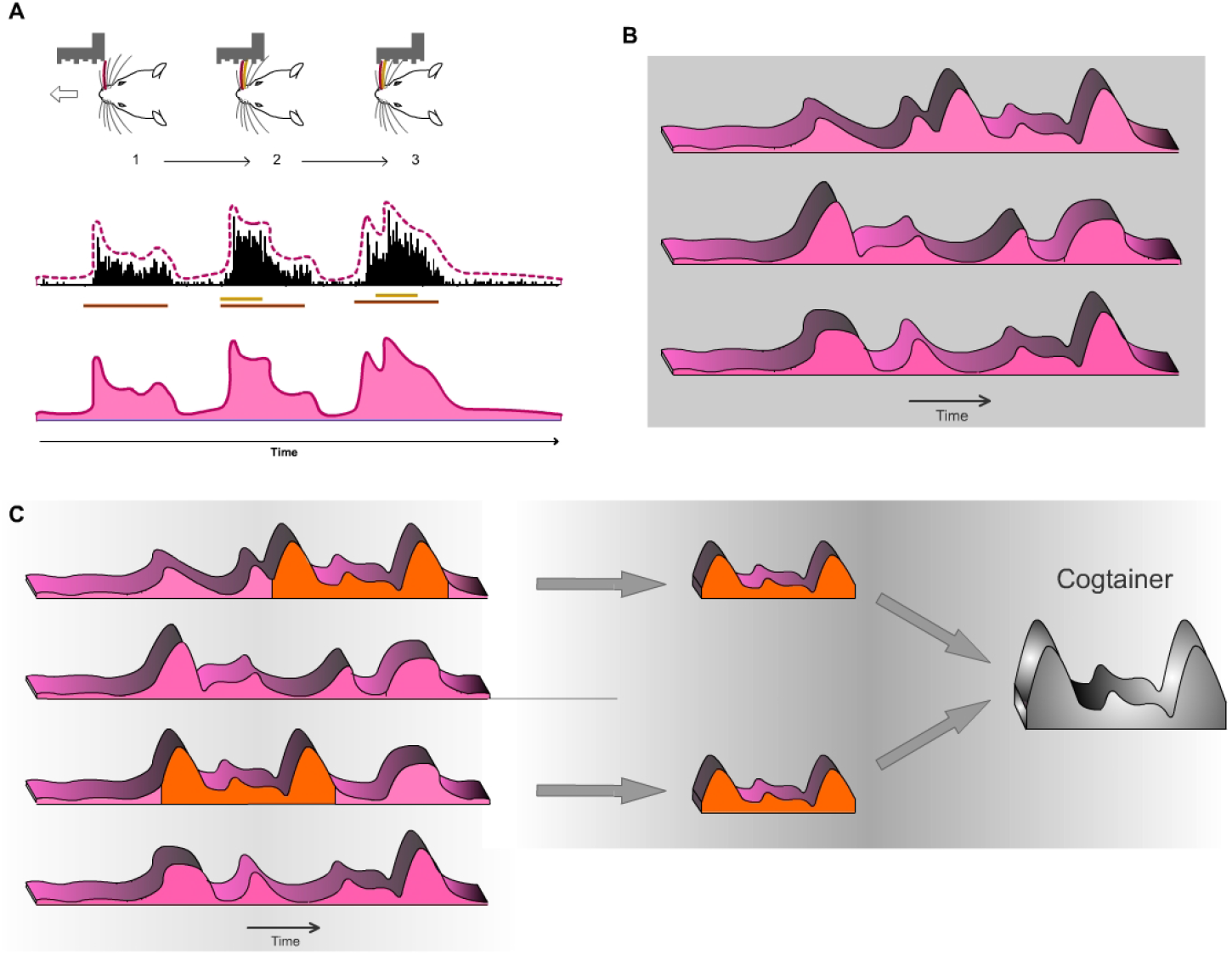
Thalamic implication in the process of extracting meaningful patterns from raw sensory information and their possible relevance in different brain functions. (A) During the tactile exploration of objects or surfaces different spatiotemporal overlappings are sequentially produced. The encoding of these overlappings by POm generates temporal sequences of activity fluctuations (1→2→3 in the example illustrated here). Functionally, the dynamical spatiotemporal structure of different sensory patterns would be accurately represented by different sustained thalamic patterns of integrated activity. Accordingly, different tactile events with diverse spatiotemporal structures would produce different ‘structured patterns of integrated activity’. Some simulated examples are shown in (B). (C) This form of encoding may provide a mechanism for detecting spatiotemporal cues, landmarks or boundaries allowing for extraction of regularities and functionally relevant patterns from the continuous flow of incoming sensory signals. By this mechanism, raw sensory input derived from the environment may be transformed into meaningful information. From this functional perspective, selected patterns of structured activity (depicted in orange) could be acquired and used as ‘Templates’ for different brain functions. Here, we propose that when these functional units are used as Templates for current or future cognitive requirements, they could be called ‘Cogtainers’.

### 6. Transmission of ‘structured patterns of integrated activity’ and its possible implication in cognition

#### Transfer of structured patterns of integrated activity across the loop

Sustained activity has been traditionally related to cognitive processes such as working memory (Fuster, 1982; Goldman-Rakic et al 1987). However, how this activity is generated, maintained and transferred is still unknown. Moreover, although interhemispheric communication during perceptual and cognitive processes is well assumed, as far as we know, no studies had been performed to define the nature of sustained activity through the thalamocortical-callosal-corticothalamic loops. These pathways have usually been studied separately. We investigated the nature and content of the activity carried by these projections and found a transmission of structured activity through this thalamocortical-callosal-corticothalamic route. Importantly, precise fluctuations in sustained activity caused by the integration of overlapping stimuli were precisely conserved (Fig. 10). Therefore, these patterns of integrated information encoded by POm were transmitted through the loop preserving their nature and their integrated structure. This finding is in agreement with our functional proposal suggesting that structured thalamic patterns of sustained activity could be transmitted and used as ‘Templates’ for different brain functions. Therefore, they must be transmitted preserving their integrity to be functionally useful.

Several important comments can be suggested. First, since POm neurons have ‘multispecific’ thalamocortical axons (Clasca et al. 2016) innervating several cortical areas including S1, S2 and M1 with different laminar profiles, our results suggest that the same ‘structured patterns of integrated activity’ generated by POm can be sent in parallel to different cortical targets. Therefore, the same message can be used by these areas and layers for different functions (i.e., perceptual, attentional and motor). Second, since POm also projects to different brain structures including the amygdala, basal ganglia, insular or ectorhinal cortex (Ohno et al. 2012), our results suggest that the same ‘Templates’ generated by POm can be used by these targets for diverse functions such as perceptual discrimination, familiarity, behavioral relevance, motivational meaning or decision-making. Accordingly, we propose that the transmission of these structured patterns of integrated activity (Cogtainers) across circuits (i.e., thalamocortical, cortico-cortical and corticothalamic circuits) supports the representation and maintenance of information during cognitive processes. From a functional perspective, this proposal provides a novel theoretical model to understand the implication of the thalamus (including POm) in cognition.

Since reciprocal interactions between the cortex (i.e., prefrontal cortex) and thalamus play a critical role in cognition, thalamic nuclei with anatomical and functional characteristics similar to those of POm may support cognitive and executive processes by the generation and transmission of Cogtainers. Indeed, thalamic responses sustaining prefrontal representations during different cognitive processes have been described (Bolkan et al., 2017; Guo et al., 2017; Schmitt et al., 2017; Rikhye et al. 2018). We propose that these effects could be the consequence of the transmission of sustained thalamic patterns of integrated activity by thalamic nuclei to their targets. The novel concept of Cogtainers could be used to functionally explain these findings.

#### POm responses to ipsilateral stimulation, POm sustained activity and its interhemispheric transmission are highly sensitive to anaesthesia

Experimental evidence has shown that during wakefulness or under light sedation, POm activity is significantly higher than during anesthetized state (Masri et al. 2008; Sobolewski et al. 2015; Zhang and Bruno 2019). This suggests that normal POm functioning can be affected in these conditions. Our findings are in agreement with this idea. In our experiments, we did not find POm responses to ipsilateral stimulation during the first hours after the application of urethane (1.3 – 1.5 g/kg i.p.). These responses were only found after sufficient time (typically 4 - 6 h) after the application of anaesthesia. Moreover, supplementary doses of urethane abolished POm responses to ipsilateral whisker stimulation (data not shown). These observations indicate that the transmission of sensory activity between hemispheres across the POm-POm loop could be highly sensitive to anaesthesia. In agreement with this, it has been previously shown that increasing the level of sedation produces the elimination of evoked responses in S1 to ipsilateral stimulation (Armstrong-James and George, 1988). This fact could explain why POm responses to ipsilateral stimulation had not been reported before. In addiction, we found that POm sustained activity was also highly affected by the level of anaesthesia (data not shown). Increasing this level by supplementary doses of urethane strongly reduced or even abolished sustained activity in POm. These findings suggest that anaesthesia reduces the interhemispheric transmission of sustained activity between cortical areas. This can have important implications in higher cognitive processes such as consciousness. Accordingly, we propose that a possible hypothesis regarding the action of anesthesia could be the suppression of thalamic sustained activity and its transmission between brain structures.

Together, this evidence indicates that high levels of sedation impair the real dynamics of POm functioning.

#### Different laminar implication in S1 in the processing of sustained activity and its transmission between hemispheres

The laminar analysis revealed that sensory-evoked responses in S1 had different temporal structures across layers (Fig. 13). We only found sustained responses lasting the duration of the stimulus in the infragranular layer. However, not all responses in the infragranular layer were sustained. This in agreement with the complexity of this layer formed by different sublayers. Sustained responses were mostly observed in the superficial part of the infragranular layer corresponding to layer 5. In addition, similar to VPM responses, supra- and granular responses were only transiently activated at the onset of the contralateral stimuli but we did not find sustained responses along stimulus presence. This is in agreement with previous findings that show higher sensory-evoked firing rates in L5 neurons (de Kock et al., 2007; Castejon et al. 2016) and sparse firing to sensory stimulation in supragranular layers of vibrissal cortex in S1 (de Kock et al., 2007; Petersen and Crochet 2013; Clancy et al. 2015; Peron et al. 2015).

In sum, our findings reveal different laminar profile of cortical responses and demonstrate different laminar implication in the processing of sustained activity and its transmission between hemispheres. This could be a common characteristic of the sensory cortex.

### 7. Differences between VPM and POm

#### Different but complementary functional roles

Important differences were observed in our study between the response modes of VPM and POm. In agreement with previous findings (Castejon et al. 2016), POm was persistently activated during whisker stimulation, whereas VPM was only transiently activated (Fig. 7A). This indicates that the effect of the stimulus duration on the response was totally different for the two nuclei. Moreover, when delivering the same spatiotemporal patterns of multiwhisker activation, we did not find a significant change of VPM responses by multiwhisker stimuli application (Fig. 7B). This is in agreement with previous findings showing that VPM response to simultaneous multiwhisker activation is very similar to individual whisker activation alone (Aguilar and Castro-Alamancos 2005). However, our results show that POm is activated more strongly by complex stimuli than by simple ones and that POm has a relevant implication in the representation of complex tactile events. Therefore, sufficient complexity is required to capture the functional dynamics of POm. Together, our results show that VPM responses are different from those of POm responses and suggest significant functional differences between POm and VPM thalamic nuclei in the processing of complex stimuli. These findings are in agreement with our hypothesis that an important function of higher-order thalamic nuclei could be the encoding of complex patterns.

In addition, our results suggest a functional difference between these thalamic nuclei underlying bilateral sensory processing. In contrast to POm, VPM did not respond to ipsilateral stimuli. Since the integration of tactile information from the two sides of the body is fundamental in bilateral perception, our results suggest a different implication of these thalamic nuclei in this function.

Taken together, these findings must be considered to functionally categorise these thalamic nuclei. The nature (structured versus discrete), type (sustained versus transient) and content (integrated versus segregated) of neural activity processed and transmitted by these nuclei may determine their functional implication and can be used to functionally classify them.

#### The hypothesis of ‘Complementary Components’

As described above, two main parallel ascending pathways convey input from the whiskers to barrel cortex (Diamond et al., 1992; Veinante et al., 2000a). This anatomical segregation suggests a different functional role of these pathways and their corresponding thalamic nuclei in somatosensory processing. Our results show that sensory stimulation protocols with similar spatiotemporal structures produce similar patterns of POm activity fluctuations even when different whiskers were activated by the same protocol. This suggests that accurate somatotopy is not a functional characteristic of this nucleus. Accordingly, we propose that to optimize the extraction of information from the sensory flow and to complement the role of POm as a general encoder of patterns, the paralemniscal system must be complemented with an additional system providing precise somatotopy. This can be the functional role of the lemniscal pathway, phylogenetically more recent and characterized by a precise somatotopy (Diamond 1995; Simons 1995).

Our results, described here, are in agreement with this proposal showing important but functionally complementary differences between POm and VPM. This functional proposal which we have called the hypothesis of ‘Complementary Components’ can explain why tactile information from whiskers is processed by parallel ascending pathways towards the cortex. This parallel architecture is also present in the majority of sensory systems in the brain and is conserved across animals (Sherman and Guillery 2006). Accordingly, we propose that sensory systems have evolved to optimize the extraction of information from the environment and that the appearance of ‘complementary’ pathways (as the somatosensory lemniscal pathway) during evolution was essential in that functional optimization.

In addition, our results demonstrate distinct laminar processing of the same stimulus by the cortex. They show that the content, type and nature of the messages that these layers receive, process and transfer is different. Therefore, different ‘Components’ are also associated with distinct laminar profiles. They may play different but complementary functional roles. This could account for the different profiles of activity in cortical layers.

## Materials and Methods

### Ethical Approval

All experimental procedures involving animals were carried out under protocols approved by the ethics committee of the Autónoma de Madrid University and the competent Spanish Government agency (PROEX175/16), in accordance with the European Community Council Directive 2010/63/UE.

### Animal procedures and electrophysiology

Experiments were performed on adult Sprague Dawley rats (220-300 g) of both sexes (40 males and 56 females). Animals were anesthetized (urethane, 1.3 – 1.5 g/kg i.p.) and placed in a Kopf stereotaxic frame. Local anaesthetic (Lidocaine 1%) was applied to all skin incisions. The skull was exposed and openings were made to allow electrode penetrations to different neuronal stations in the trigeminal complex, thalamus and cortex.

Our recordings were mostly performed several hours after the application of urethane (typically after 5 - 6 h). To assure the absence of whisker movements and pinch withdrawal reflexes, supplementary dosis of urethane were applied if necessary.

Extracellular recordings were made in the Principal (PrV; Posterior from bregma 9–10; Lateral from midline 3–3.5, Depth 8.5–9.5; in mm) and Interpolar trigeminal nuclei (SpVi; P 11.5–14; L 2.5–3.5, D 8.5–9.5) of the trigeminal complex, in the posteromedial thalamic nucleus (POm; P 2.5-4.5, L 2-2.5, D 5-6.5), in the ventral posteromedial thalamic nucleus (VPM; P 2.8-4.6, L 2-3.5, D 5.5-7) and in the vibrissal region of the primary somatosensory cortex (S1; AP 0.5-4, L 5-7). Laminar recordings in supra- (D 150 – 550 μm), granular (D 650 – 850 μm) and infragranular (D > 950 μm) layers of S1 were also performed. Unanalyzed gaps were left between layers to compensate for differences in cortical thickness across this area and as a safeguard against potential errors in laminar localization. Tungsten microelectrodes (2–5 MΩ) were driven using an electronically controlled microdrive system (DavidKopf).

### Sensory stimulation and patterns generation

Sensory stimulation was characterized by spatiotemporal patterns of multiwhisker deflections simulating possible real complex stimuli or sequences of stimuli similar to those occurring in natural circumstances. Details of the multiwhisker stimulation patterns are described in Fig 2A, Fig 3A, Fig. 4A and Fig. 5A, B. Anesthetized rats were used to facilitate their application. Using a pneumatic pressure pump (Picospritzer) that delivers air pulses through polyethylene tubes (1 mm inner diameter; 1-2 kg/cm^2^), sensory patterns were generated using controlled multiwhisker deflections performed by overlapping air puffs of different durations (20-2000 ms) applied to different whiskers in one or both sides of the face and avoiding skin stimulation. Accordingly, many whiskers were activated simultaneously producing different spatiotemporal overlapping dynamics. The air-puffers were precisely placed and the whiskers were trimmed to a length of 10–30 mm to allow precise overlapping stimulations. Overlappings produced by the activation of whiskers in different directions were also included. A variant order was adopted for delivering the stimulation patterns to avoid possible temporal dependency. We applied 20-70 trials per pattern at low frequency (0.3 - 0.5 Hz). Receptive field sizes were determined by deflecting individual vibrissae with a hand-held probe and monitoring the audio conversion of the amplified activity signal.

### Inactivation and lesion of thalamic nuclei and cortical areas

We inactivated the cortex with local infusions of lidocaine (10%) or muscimol (1 mg/ml). To further evaluate the proper level of cortical inactivation, the basal activity and sensory responses to whiskers activation were continuously checked in the inactivated cortical area. Infusions were repeated every 15 min until cortical activity recorded in deep layers was silenced, on average 25 min after the first application. Pharmacological deactivation of POm was performed by injecting 100-200 nL of muscimol (1 mg/ml) in this thalamic nucleus. The drug was slowly delivered through a cannula connected to a Hamilton syringe (1μl) over a one-minute period.

Lesions of different cortical areas were also performed in our experiments. To assure the precision of cortical lesions, the skull was exposed and openings were precisely restricted to the corresponding cortical area according to stereotaxic coordinates (S1, described above; secondary somatosensory cortex (S2), P 0–3.7; L 5.5 – 7.5; primary motor cortex (M1), A 0.5 – 2.5, L 0.2 – 3). Lesions were made by cutting and aspirating the cortical tissue and included superficial and deep layers of these areas.

### Histology

After the last recording session, animals were deeply anesthetized with sodium-pentobarbital (50 mg/kg i.p.) and then perfused transcardially with saline followed by formaldehyde solution (4%). After perfusion, brains were removed and postfixed. Serial 50 μm-thick coronal sections were cut on a freezing microtome (Leica, Germany). These sections were then prepared for Nissl staining histochemistry for verification of electrodes tracks, delimitation of cortical lesions and discrimination of thalamic nuclei. Positions of the electrode tips and extensions of cortical lesions were histologically verified by comparing these coronal brain sections with reference planes of the rat brain stereotaxic atlas (Paxinos and Watson 2007).

### Data acquisition and analysis

Data were recorded from PrV, SpVi, VPM, POm and S1. The raw signal of the *in vivo* extracellular recordings was filtered (0.3–5 kHz), amplified (DAM80 preamplifier, WPI) and digitalized (Power 1401 data acquisition unit, CED). Single-unit activity was extracted with the aid of Spike2 software (CED) for spike waveform identification and analysis. Multi-units were collected by amplitude sorting. We defined response magnitude as the total number of spikes per stimulus occurring between response onset and offset from the peristimulus time histogram (PSTH, bin width 1 ms unless noted otherwise). Response onset was defined as the first of three consecutive bins displaying significant activity (three times higher than the mean spontaneous activity) after stimulus and response offset as the last bin of the last three consecutive bins displaying significant activity. Response duration was defined as the time elapsed from the onset to offset responses. In all figures, raster plots represent each spike as a dot and each line corresponds to one trial. Spikes were aligned on stimulus presentation (Time 0 ms). All data are expressed as the mean ± standard error of the mean (SEM). Error bars in the figures correspond to SEM. For normally distributed data (Shapiro-Wilk normality test), statistical analyses were conducted using a Student’s t-test. Non-normally distributed data were compared with the Wilcoxon matched-pairs test. Multiple comparisons were evaluated using a One-way ANOVA test. Statistical significance was considered at *P < 0.05, **P < 0.01, ***P < 0.001.

## Additional information

### Data availability statement

The data that support the findings of this study are available from the corresponding author upon reasonable request.

### Author contributions

C.C. conceived the theoretical models and hypotheses, designed and conducted the experiments, analyzed the results and wrote the paper.

J.M.C. conducted the experiments, analyzed the results and reviewed the paper.

A.N. designed and conducted the experiments, analyzed the results, reviewed and edited the paper.

